# Methanotrophic *Methanoperedens* archaea host diverse and interacting extrachromosomal elements

**DOI:** 10.1101/2023.08.02.551345

**Authors:** Ling-Dong Shi, Jacob West-Roberts, Marie C. Schoelmerich, Petar I. Penev, LinXing Chen, Yuki Amano, Shufei Lei, Rohan Sachdeva, Jillian F. Banfield

**Affiliations:** Innovative Genomics Institute, University of California, Berkeley, CA, USA; Environmental Science, Policy and Management, University of California, Berkeley, CA, USA; Sector of Decommissioning and Radioactive Wastes Management, Japan Atomic Energy Agency, Ibaraki, Japan; Earth and Planetary Science, University of California, Berkeley, CA, USA

## Abstract

Methane emissions that contribute to climate change can be mitigated by anaerobic methane-oxidizing archaea such as *Methanoperedens*. Some *Methanoperedens* have huge extrachromosomal genetic elements (ECEs) called Borgs that may modulate their activity, yet the broader diversity of *Methanoperedens* ECEs is little studied. Here, we report small enigmatic linear ECEs, circular viruses and unclassified ECEs, that we predict replicate within *Methanoperedens*. The linear ECEs have features such as inverted terminal repeats, pervasive tandem repeats, and coding patterns that are strongly reminiscent of Borgs, but they are only 52 kb to 145 kb in length. They share proteins with Borgs and *Methanoperedens*. Thus, we refer to them as mini-Borgs. Mini-Borgs are genetically diverse and we assign them to at least five family-level groups. We also identify eight novel families of *Methanoperedens* viruses, some of which encode multiheme cytochromes, and unclassified circular ECEs that encode TnpB genes. A population-heterogeneous CRISPR array is in close proximity to the TnpB and has spacers that target other *Methanoperedens* ECEs including previously reported plasmids. The diverse groups of ECEs exchange genetic information with each other and with *Methanoperedens*, likely impacting the activity and evolution of these environmentally important archaea.

## Introduction

Methane (CH_4_) is the second most important greenhouse gas and contributes substantially to global climate change. Methane is generated by methanogenic archaea and can be consumed by methanotrophic bacteria and archaea^1,2^. Anaerobic methanotrophs (ANME) that perform anaerobic oxidation of methane (AOM) form a polyphyletic group comprised of ANME-1^3^, ANME-2a-c^4^, ANME-3^5^, and ANME-2d (also known as *Methanoperedens*)^6^. Unlike most ANME that inhabit marine sediments and rely on syntrophic partners, *Methanoperedens* usually live in freshwater environments and independently couple the reduction of nitrate^7^, iron/manganese oxides^8,9^, and other potential extracellular electron acceptors^10,11^, to methane oxidation. It was recently proposed that the activity of some *Methanoperedens* species can be modulated by extrachromosomal genetic elements (ECEs), including extraordinarily large, linear Borgs (up to ∼1.1 Mbp) and circular plasmids (up to ∼253 kb)^12,13^. The Borg genomes are particularly notable in that they encode diverse, and in some cases expressed^14^, metabolism-relevant genes, including methyl-coenzyme M reductase (MCR) and multiheme cytochromes (MHCs), potentially linking their activity to methane oxidation rates. Echoing this, phages of aerobic methane-oxidizing bacteria carry a subunit of methane monooxygenase that may impact host bacterial methane oxidation rates^15^. However, little is known about *Methanoperedens* viruses, although the presence of proviruses has been reported^16^. This raises the possibility that *Methanoperedens*-associated viruses and other ECEs may also modulate host abundance and methane oxidation activity in terrestrial environments linked to methane emissions.

To explore the diversity of ECEs associated with *Methanoperedens*, we investigated wetland soils known to contain *Methanoperedens* and Borgs and found diverse novel ECEs that we predict replicate in *Methanoperedens*. Related sequences were also recruited from deep terrestrial subsurface sedimentary rocks and public databases. Most of the ECE genomes were manually curated to completion, enabling confident analyses of their genome architecture and genetic repertoires.

## Results

### Mini-Borgs are new ECEs associated with *Methanoperedens*

By analysis of assembled metagenomic data from previously studied deep, *Methanoperedens*-containing wetland soil in Lake County, CA^12^, we identified taxonomically unclassified low GC (31.6% to 35.9%) contigs that primarily encode hypothetical proteins as potential genomic fragments of ECEs (Fig. 1a). Eight sequences were manually curated to completion and found to represent linear genomes of 52,459 bp - 106,520 bp in length, terminated by inverted repeats (Fig. 1b). Most have two replichores of very unequal lengths, with genes on a single strand of each replichore, as observed in Borgs^17^. However, the small replichore is proportionally slightly shorter than that in Borgs, and some have only a single replichore (mostly when lengths are <60 kb). As for Borgs^12^, these ECEs are predicted to initiate replication from the termini, based on the cumulative GC skew patterns (Fig. S1).

**Fig. 1.**
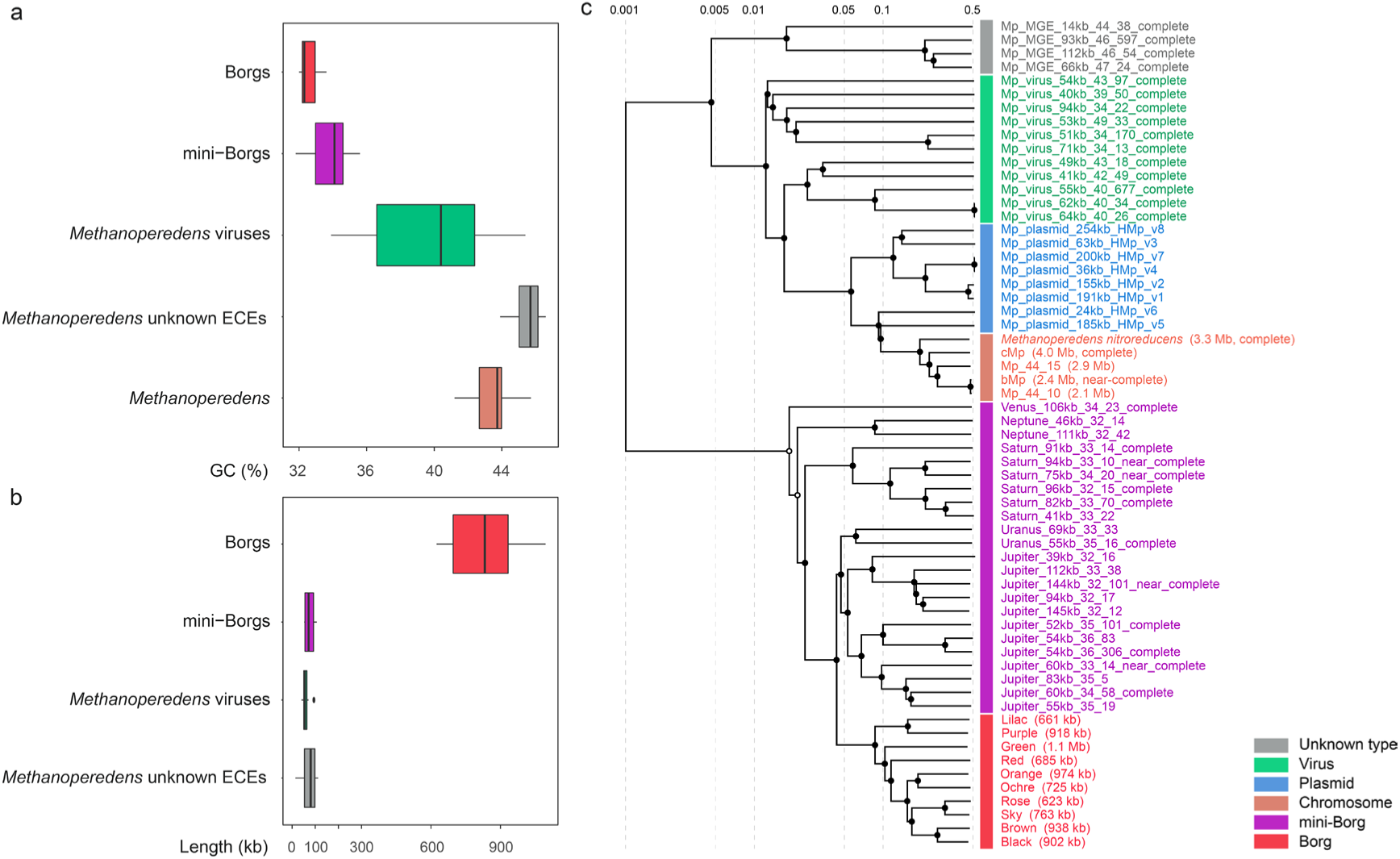
Comparison of genomic features of *Methanoperedens* and associated ECEs. **A**. GC content of complete Borgs (n=10), mini-Borgs (n=8), *Methanoperedens* viruses (n=11), and unknown types of ECEs (n=4), compared to *Methanoperedens* genomes. The black lines inside the boxes indicate the median, and the box edges show the interquartile range. **B.** Length ranges of ECEs that associate with *Methanoperedens*. **C**. Global proteome-based similarity of *Methanoperedens* and associated ECEs. Bar colors indicate *Methanoperedens* and ECE types. Mini-Borgs clades are defined as similarity scores ≥ 0.05 and names.

Three related ECE sequences have a tiny second replichore (Fig. S2) but could not be curated into the terminal repeats, and a fourth one is terminated by perfect inverted repeats but has a gap that is spanned by paired reads (Fig. S3). These ECE sequences display a strong and symmetrical decrease in read coverage towards the genome ends (Fig. S2). As the replichore structure precludes the explanation due to rapid replication initiated from the center of the genome^18^, we attribute this coverage pattern to DNA degradation of the unprotected linear sequence ends. Similar degradation patterns have been documented for linear mitochondrial DNA^19^. We infer that DNA degradation impeded the completion of these four genomes. Given the average paired read insert size, we estimate the full length of the fourth ECE sequence is ∼145 kb. Further, we identified 11 more partial genomes, bringing the total number of the new type of related ECE sequences to 23 (Table S1).

A typical genomic feature of the ECE sequences is pervasive perfect tandem repeat (TR) regions, which locate both within and between genes (Table S1 and Fig. S4). TR within genes have unit repeat lengths divisible by three, thus introducing amino acid tandem repeats. In one case, a TR region is located within the inverted terminal repeats (ITRs). It has been suggested that intergenic TR may function as regulatory RNAs^17^, and those in the ITRs may regulate replication from the genome ends.

Of the predicted proteins encoded in these ECE genomes, ∼85% have no functional prediction based on the UniProt and KEGG databases^20,21^. Functionally annotated proteins are mainly involved in the transfer of glycosyl and methyl groups, as well as in energy conservation, such as archaeal-type ATP synthase subunit K. ATP synthase subunit K sequences are also present in Borgs and *Methanoperedens* but the protein sequences form independent sibling clades. The tree topology suggests that these ECEs and Borgs acquired this gene from a common ancestor likely related to *Methanoperedens* (Fig. 2a). Similar phylogenetic topologies are observed for replication factor C small subunit (Fig. S5), myo-inositol-1-phosphate synthase (Fig. S6), MBL (metallo-β-lactamase) fold metallo-hydrolase (Fig. S7), DNA helicase RuvB (Fig. S8), and a hypothetical protein (Fig. S9). In combination, these observations indicate that the newly described ECEs are related to both Borgs and *Methanoperedens*.

**Fig. 2.**
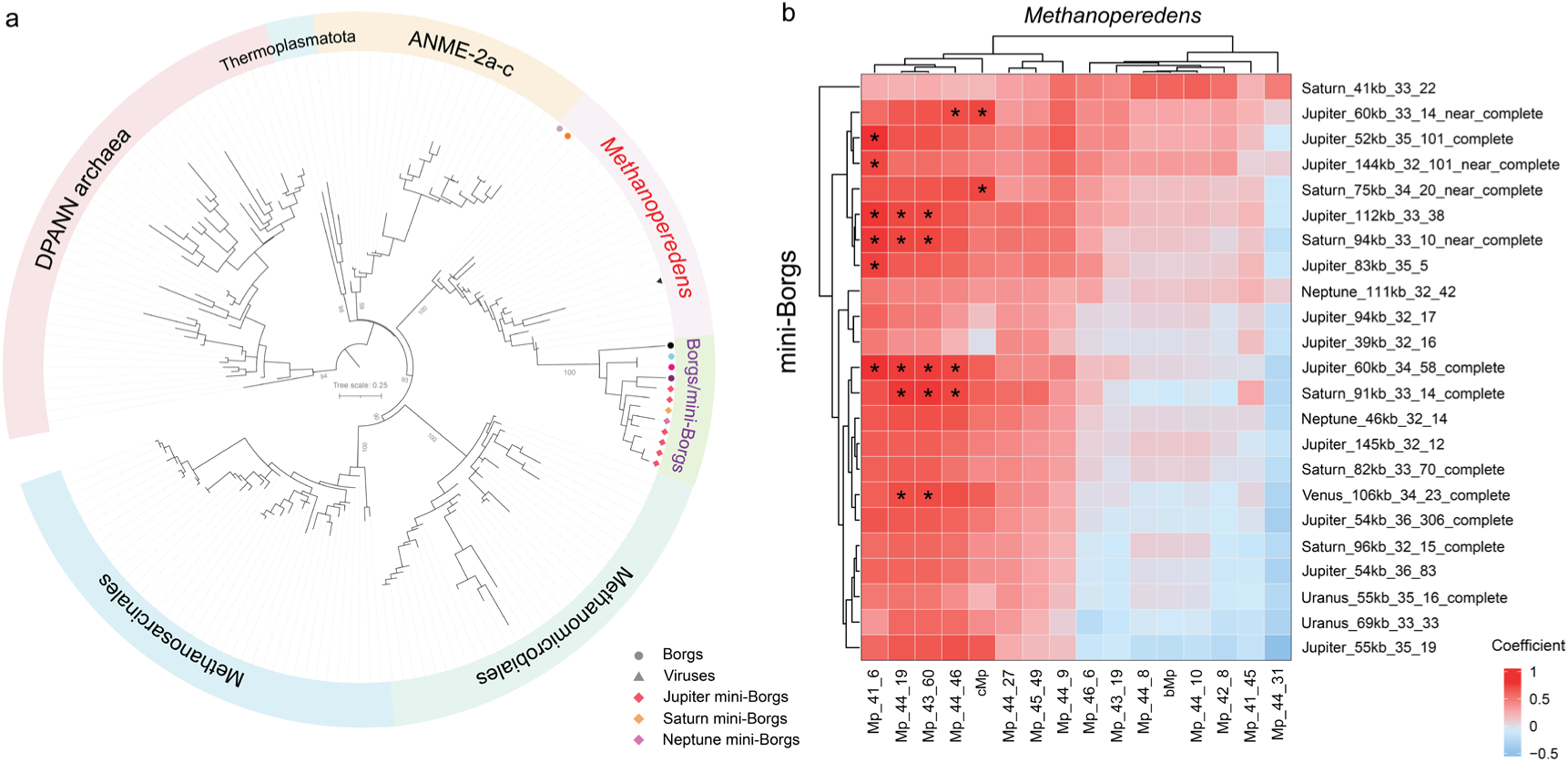
Evidence linking mini-Borgs to *Methanoperedens*. **A.** Phylogenetic tree of archaeal A-type ATP synthase subunit K (AtpK). Reference sequences were downloaded from NCBI. Arc colors indicate different taxonomic clades. The tree was mid-point rerooted and support values were calculated based on 1000 replicates. **B.** Heatmap of abundance correlations between mini-Borgs and *Methanoperedens*. Abundances were calculated across 68 wetland samples. Colors indicate Spearman correlation coefficients and stars denote high positive correlations (i.e., *ρ* ≥ 0.7 and *p* ≤ 1×10^−10^).

Extremely long branch lengths for the ECE groups compared to *Methanoperedens* may suggest rapid evolution of these ECEs. Given the phylogenetic information, the presence of pervasive genic and intergenic TRs, and overall genome architectures similar to those of Borgs (Fig. 1a), we infer that these newly described ECEs are related to Borgs. In view of their much smaller genomes (averagely ∼10 times smaller) (Fig. 1b), we refer to them as “mini-Borgs”.

Mini-Borgs lack a near-universal phylogenetically informative protein, such as the ribosomal protein L11 (rpL11) that is used to classify Borgs. Thus, we used global proteome similarity to define clades, and named the clades after planets (Fig. 1c). The Borg proteomes cluster together in a clade that is sibling to the Jupiter and Uranus mini-Borg clades (Fig. 1c). Mini-Borgs are proteomically more diverse than Borgs. Of the 40 essentially syntenous genes that are distributed throughout the large replichore of Borgs (likely inherited from a common ancestor)^14^, two occur in mini-Borgs. However, these two do not occur outside of Borgs and mini-Borgs (Fig. S10). As these marker proteins individually define similar topological relationships, we concatenated them to better resolve the relationship between Borgs and mini-Borgs. The resulting phylogenetic tree indicates that they are evolutionarily related but distinct groups (Fig. S10).

As mini-Borgs lack anything approaching the complete machinery required for replication, it is clear that they require a host organism. Given phylogenetic trees pointing to gene acquisition from *Methanoperedens*, we infer that the mini-Borgs replicate in *Methanoperedens*. In many samples, the mini-Borgs are either at low coverage or undetected. However, we were surprised to note the extremely high relative abundances of two mini-Borgs compared to any potential coexisting host organisms. For example, a 60 cm deep soil sample has a mini-Borg (Jupiter_52kb_35_101_complete) with a coverage of > 8000 x, and the most abundant organism, a Bathyarchaeota, has a coverage of ∼130 x. This implies a genome copy ratio of > 60 : 1, and the copy number would be far greater if *Methanoperedens* species, individually (i.e., > 600 : 1) or in combination, are the hosts.

As the gene phylogenies strongly suggest a dependence of mini-Borgs on *Methanoperedens* rather than Bathyarchaeota, we statistically compared the abundances of mini-Borgs and *Methanoperedens* across 68 wetland soil samples (collected from the surface to a depth of 1.75 m) to test for specific mini-Borg - host linkages based on co-occurrence. This analysis shows significant positive correlations between certain mini-Borgs and *Methanoperedens*, and a combination of several possible hosts for each mini-Borg (Fig. 2b). One *Methanoperedens* species could host only one mini-Borg (e.g., Mp_44_31) or host multiple different mini-Borgs (e.g., Mp_43_60). In line with the latter case, we detected evidence for recombination among mini-Borgs, which requires their co-existence in the same host. For example, Jupiter_54kb_36_306_complete has one genomic region where mapped reads clearly show a variety of linkage patterns for distinct nucleotide polymorphism motifs (Fig. S11).

### *Methanoperedens* viruses may augment electron transfer

In addition to mini-Borgs, we identified ECE fragments classified as viral based on the presence of structural genes. A subset of these viral sequences could be circularized and curated to completion. We predict that these viruses associate with *Methanoperedens* based on sequence similarity and CRISPR spacer targeting (Table S2). Related viral sequences were identified in the deep terrestrial subsurface sedimentary rocks^22^ and the public IMG/VR v4 database^23^. In total, 11 distinct *Methanoperedens* viruses were identified. These viruses generally have GC contents comparable with those of the predicted host *Methanoperedens*, and genome lengths ranging from ∼40 kb to ∼94 kb (Fig. 1). The putative viruses share few genes and do not cluster with known prokaryotic viruses/phages (Fig. 3a).

**Fig. 3.**
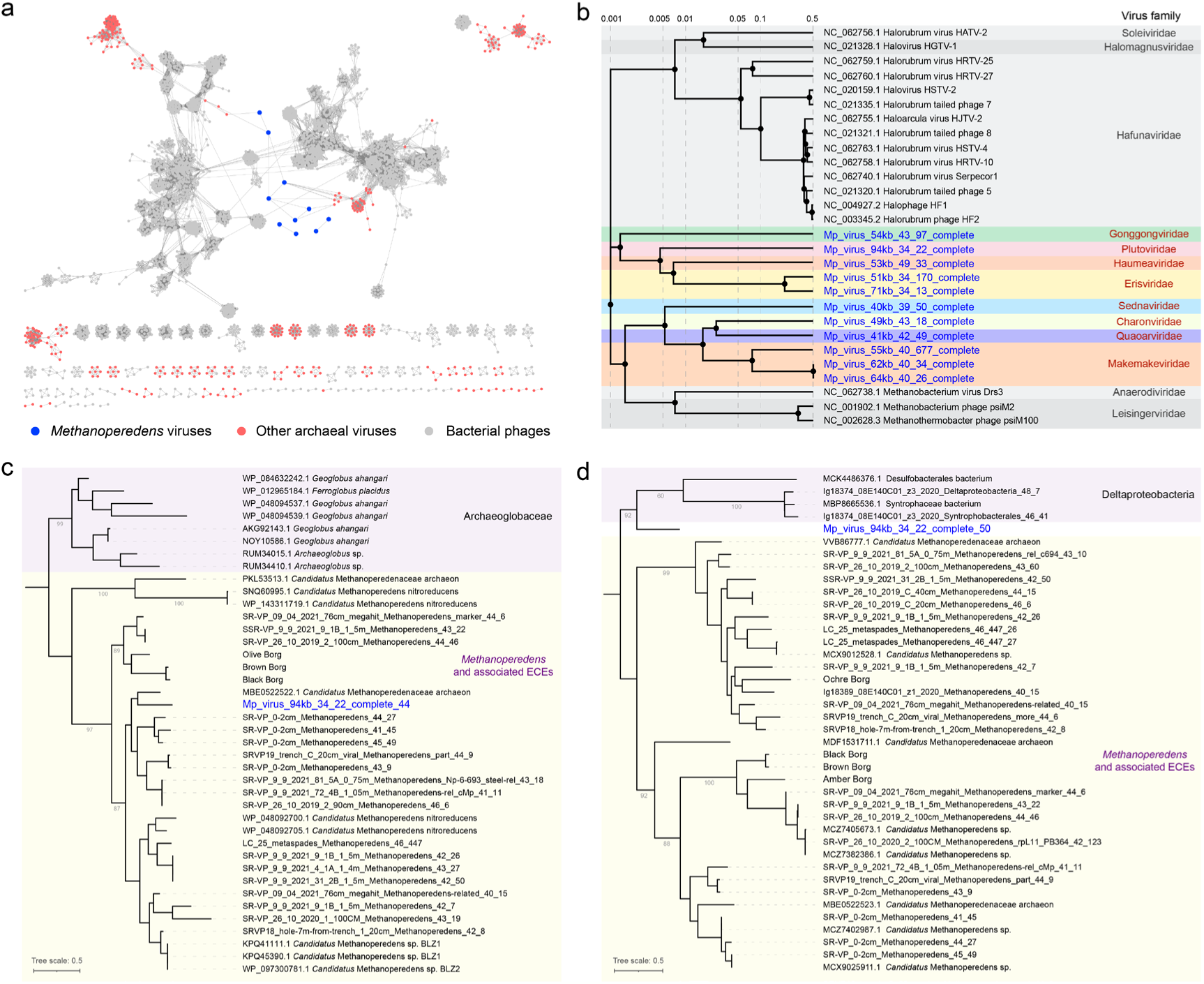
Genomic comparison and phylogeny of *Methanoperedens* viruses. **A**. Gene-sharing network for *Methanoperedens* viruses and RefSeq prokaryotic viral genomes. Nodes indicate viral genomes and edges indicate shared gene content. One virus classified as a singleton is not included in the network. **B.** Global proteome-based phylogenetic analyses of *Methanoperedens* viruses and other archaeal viruses. Background colors distinguish different viral families. **C.** Phylogenetic trees for multiheme cytochromes No. 44 (five binding sites) and **D.** No. 50 (six binding sites). Blue text indicates the proteins from *Methanoperedens* viruses. The tree was mid-point rerooted and support values were calculated based on 1000 replicates.

A global proteome-based phylogenetic analysis demonstrated that the *Methanoperedens* viruses could be assigned to approximately eight families (Fig. 3b). This was confirmed by genome synteny, where viruses from the same family have more syntenous gene homologs than those belonging to different families (Fig. S12). We tentatively assign the *Methanoperedens* viruses to Plutoviridae, Erisviridae, Makemakeviridae, Gonggongviridae, Haumeaviridae, Charonviridae, Quaoarviridae, and Sednaviridae, referencing the asteroids in the solar system.

*Methanoperedens* viruses have some interesting genes. Two viral genomes (i.e., Mp_virus_94kb_34_22_complete and Mp_virus_71kb_34_13_complete) encode a single Cas4-like protein which forms a separate clade from Cas4 in coexisting *Methanoperedens* (Fig. S13). These viruses likely obtained Cas4-like genes from *Methanobrevibacter,* based on the close amino acid similarity to the Cas4 of these archaea. In CRISPR systems, Cas4 proteins often coexist with Cas1 and Cas2 and contribute to the incorporation of new spacers into the CRISPR locus^24^. However, isolated, CRISPR-independent virus-encoded Cas4 proteins may confer CRISPR-Cas interference activity by misleading the defense system, for example by limiting spacer acquisition or by triggering incorporation of erroneous spacers of host origin^25,26^. This suggests an anti-CRISPR role of Cas4-like proteins encoded in *Methanoperedens* viruses.

One virus, Mp_virus_41kb_42_49_complete, encodes a type IV-B Cas system consisting of *csf1* (*cas8*), *cas11*, *csf2* (*cas7*), *csf3* (*cas5*), and a non-canonical *cysH* gene, but with no CRISPR array nearby. Phylogenetic analysis of the Csf1 protein suggests a bacterial origin (Fig. S14), possibly from *Candidatus* Omnitrophica which has been predicted to have a syntrophic relationship with *Methanoperedens* in a deep granitic environment^27^. Although type IV-B systems are widely distributed in bacteria and archaea and common in plasmids^28^, homologs have previously not been identified in *Methanoperedens*. Based on structure analysis, the type IV-B complex was proposed to inactivate small guide RNAs (e.g., crRNAs), enabling plasmids/viruses to evade CRISPR targeting by their hosts^29,30^. Given the type IV-B Cas system encoded in the genome, it is conceivable that *Methanoperedens* viruses employ that to mitigate the activity of their host’s defense system.

In addition to genes involved in viral survival, *Methanoperedens* viruses encode several metabolic genes that may contribute to host activity. Most conspicuous are multiheme cytochromes (MHCs). MHCs are common in archaea and bacteria where they play roles in respiratory complexes (e.g., nitrate reductase, hydroxylamine oxidoreductase) and disposal of electrons to external electron acceptors^31,32^. They have been reported in all well-sampled Borg genomes^14^ but have never been found in viruses/phages. One archaeal virus predicted to infect *Methanoperedens* (Mp_virus_94kb_34_22_complete) encodes two extracellular MHCs with five and six heme-binding sites. One (No. 44), with five heme-binding motifs, clusters with homologs of *Methanoperedens* and associated Borgs (Fig. 3c). The other one (No. 50) is relatively closely related to those of *Methanoperedens* and Deltaproteobacteria, but distinct from both (Fig. 3d). Alignment of viral and microbial MHCs, including a homolog of *Archaeoglobus veneficus* that is experimentally proven capable of long-range electron transfer^33^, shows highly conserved His and Cys residues (Fig. S15). This suggests that virus-borne MHCs have the potential for extracellular electron transfer and may augment the metabolic activity of the host *Methanoperedens* during infection.

### Group II introns shared by *Methanoperedens* and their viruses

Two *Methanoperedens* viruses belonging to Makemakeviridae are essentially identical except for a ∼2 kb region (Fig. S16a). Read mapping showed much lower coverage over that region compared to the flanking genome, indicating that this region is in only a subset of the virus genomes (Fig. S16b). Supporting this, some paired reads perfectly span the ∼2 kb region. The region encodes a group II intron reverse transcriptase that is highly similar to those encoded by *Methanoperedens* and clusters with them, suggesting recent gene transfer (Fig. S17). Further, there is a surprisingly high similarity between the viral ∼2 kb region and those in genomes of coexisting *Methanoperedens*, suggesting that not only the reverse transcriptase, but the whole region originated from *Methanoperedens* (Fig. S18).

In addition to the reverse transcriptase, the conserved ∼2 kb regions also harbor genes of unknown functions (Fig. S18). For example, the region in the *Methanoperedens* genome SR-VP_Mp_41_11 carries two genes whereas the region in LC_25_Mp_44_558 carries one. Such an observation is exceptional, as introns are normally considered to move only themselves^34^. Our finding suggests that the group II intron may contribute to gene transfer between *Methanoperedens* and its associated viruses, similar to transposons that transfer, for example, antibiotic resistance genes^35^.

### Unknown ECEs with TnpB-associated CRISPR arrays target other *Methanoperedens* ECEs

In addition to the mini-Borgs and viruses, we identified four unclassified ECEs that appear to associate with *Methanoperedens* based on CRISPR- and blast-based predictions (Table S3). They have similar GC contents to *Methanoperedens* and comparable genome sizes to mini-Borgs and viruses (Fig. 1a-b). Clustering based on the composition of the proteomes shows they are more genomically analogous to *Methanoperedens*-associated viruses and plasmids than to mini-Borgs or Borgs. However, although circularized and complete, neither viral nor plasmid hallmark genes were detected (Fig. 1c).

We identified transposon-associated TnpBs adjacent to perfect CRISPR arrays in two of these ECEs. In the representative genome Mp_ECE_93kb_46_597_complete, two CRISPR arrays with four and eight repeats have putative TnpBs within two genes (Fig. 4a). Structure modeling by AlphaFold2^36^ indicates that the TnpB adjacent to the larger CRISPR array adopts a conformation structurally homologous with Cas12. The predicted protein has the conserved binding sites for DNA recognition and cleavage (Fig. 4b), suggesting potential endonuclease activity. Intriguingly, the adjacent CRISPR array has a highly diverse spacer content and variable locus length, suggesting that it is functionally active (Fig. 4c).

**Fig. 4.**
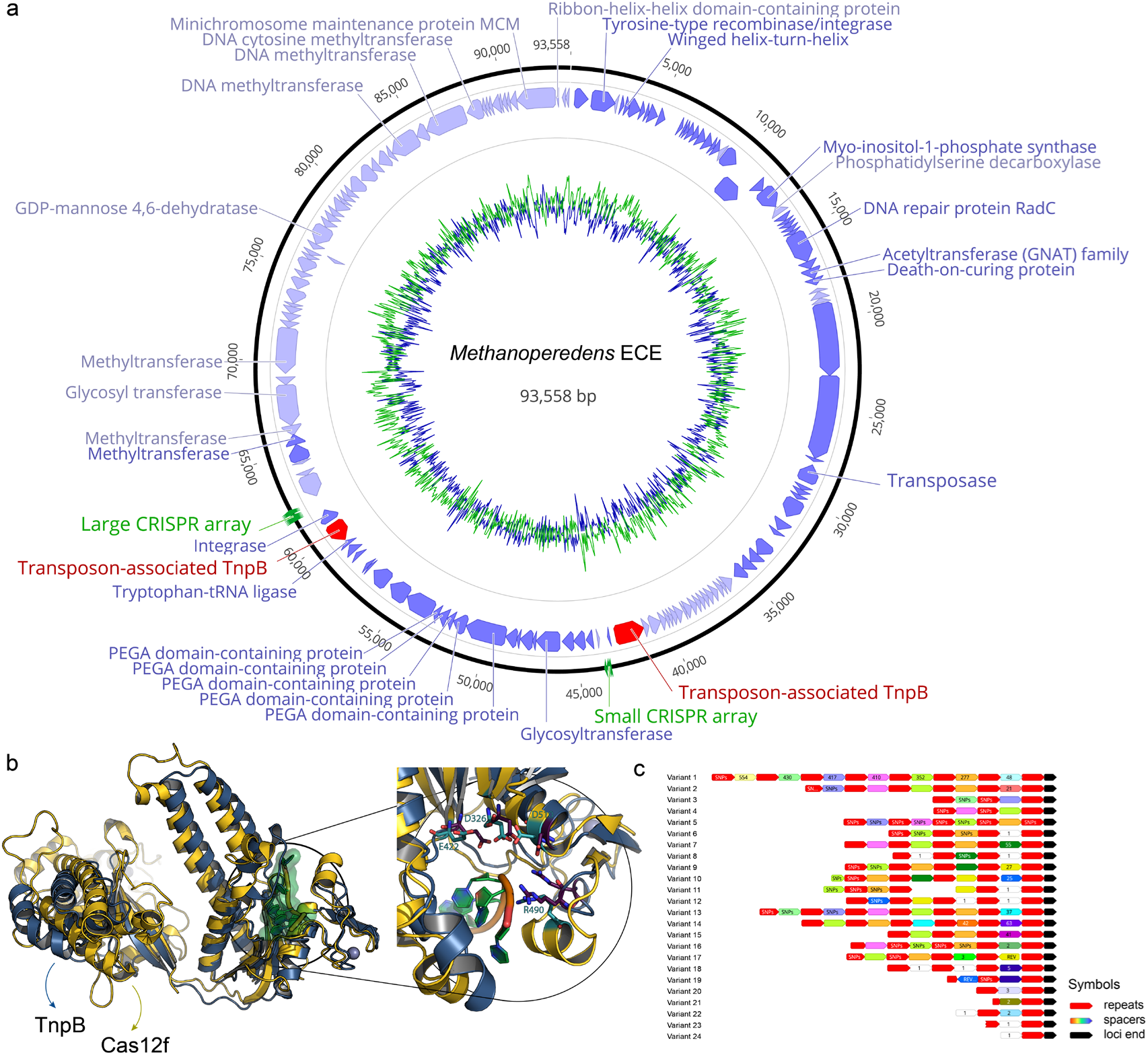
TnpB and associated CRISPR array in an unclassified circular ECE. **A.** Genome of Mp_ECE_93kb_46_597_complete. The inner green and blue circles indicate AT and GC content. Arrows indicate genes. **B.** Structural superimposition for the TnpB model (blue) and cryo-EM structure of Cas12f (yellow; RCSB PDB ID: 7l49). The target DNA where the cleavage is performed is shown by the green surface. A detailed view of the active site in the nuclease domain RuvC is shown. Active site residues from 7l49 are highlighted with cyan and corresponding residues in the TnpB model are purple. **C.** Extensive spacer diversity towards the end of the large TnpB-associated CRISPR locus. Spacers with different colors have different sequences, except for white spacers, each of which is novel. Numbers inside spacers indicate how often the spacer was found in the reads.

Deep sampling of the heterogeneous CRISPR locus enabled a large inventory of 1,527 unique spacer sequences. We used these to identify the sequences targeted by the TnpB-associated CRISPR. One target is a transposase belonging to the IS1634 family that is encoded in both *Methanoperedens* chromosomes and an associated plasmid (Fig. S19). The targeted transposases are dissimilar from those of other microorganisms (Fig. S20).

Another spacer targets an intergenic region in a circularized but unclassified element, Mp_ECE_14kb_44_38_complete (Figs. 1c and S19), for which we identified no viral or plasmid marker genes. Given the host for this ECE is predicted to be *Methanoperedens* based on genome similarity, we conclude that the TnpB-associated CRISPR harbored by *Methanoperedens* ECEs can target other mobile elements, thus protecting the host against infection.

### Gene transfer between *Methanoperedens* and divergent ECEs

Given the large variety of ECEs that we now predict to associate with *Methanoperedens* and that co-occur with this archaeon in the wetland soil, we investigated the evidence for lateral gene transfer amongst them. Of particular interest were the TnpB genes, which are encoded by *Methanoperedens* and all of its associated ECEs (Fig. S21). The TnpBs appear to be derived from a variety of sources, and some in Borgs are phylogenetically closer to those in Asgard archaea than in *Methanoperedens*. Other TnpBs in the ECEs form three distinct clades, all of which include corresponding homologs from *Methanoperedens* (Fig. S21). Such tight phylogenetic affiliation supports the predicted host-ECE relationships and provides evidence for genetic exchange. The largest clade includes members from *Methanoperedens*, Borgs, viruses, plasmids, and unknown ECEs and is located nearest the ANME-2a/2b cluster. The organismal topology is congruent with the genome phylogeny of archaea^37^, suggesting a vertical gene transfer of TnpB from a common ancestor into ANME and then to their ECEs (Fig. S21). However, another clade containing *Methanoperedens*, Borgs, and one mini-Borg adjoins the *Methanosarcina* cluster, distant from the ANME-2 group. This suggests that *Methanoperedens* (or their ECEs) may have acquired the TnpB from *Methanosarcina.* A similar scenario may explain the clade comprised of *Methanoperedens* and unclassified ECEs that is adjacent to a *Methanothrix* cluster (Fig. S21).

Besides TnpB, *Methanoperedens* harbor a transposase that is shared with its ECEs. Based on phylogeny, this transposase has moved between *Methanoperedens*, Borgs, viruses, and plasmids (Fig. S22). Also of interest is a thymidylate synthase gene (*thyA*) (Fig. S23) and a gene encoding a hypothetical protein (Fig. S24) that are shared by *Methanoperedens* and its ECEs. Thymidylate synthase methylates deoxyuridine monophosphate (dUMP) into deoxythymidine monophosphate (dTMP) for the production of thymidine, an essential precursor for DNA synthesis^38^. Possession by Borgs, viruses, and plasmids may facilitate *Methanoperedens* thymidine synthesis thus enhancing their viability. The latter shared but unclassified protein is only found in *Methanoperedens*, Borgs, viruses, and plasmids, strongly suggesting the gene transfer among them. The last shared gene currently identified encodes DNA repair endonuclease (ERCC4) and occurs in *Methanoperedens*, viruses, and plasmids but not in Borgs or mini-Borgs (Fig. S25). This enzyme is responsible for repairing DNA damage and maintaining genome stability^39^. Transferring the gene to varied ECEs may enhance the survival of *Methanoperedens*. Overall, our phylogenetic analyses suggest extensive lateral gene transfer among *Methanoperedens* and its associated ECEs.

## Discussion

Extrachromosomal elements are integral components of biological systems and impact organism abundances, activity levels, genetic repertoires, and evolutionary trajectories. In the case of anaerobic methane-oxidizing *Methanoperedens* archaea, there is now evidence that unusual Borg ECEs and some plasmids carry a variety of interesting genes and replicate within *Methanoperedens* cells^12,13^. But are these part of a larger constellation of ECEs with the potential to impact *Methanoperedens* activity *in situ*? Here we discover and genomically describe mini-Borgs, viruses, and other unclassified ECEs, and provide evidence that they form an interconnected, gene-sharing network. Through their interactions with each other and *Methanoperedens*, these ECEs may contribute to the proclivity of this archaeon for gene acquisition via lateral transfer^10^, and therefore modulate the metabolic activity of *Methanoperedens*.

A striking observation is that mini-Borgs exhibit a vast range in relative abundances compared to the abundances of coexisting organisms. In the most extreme case, the abundance ratio of one mini-Borg to the most abundant organism (a Bathyarchaeota) is > 60 : 1. At the DNA level, this would imply similar amounts of mini-Borg and archaeal DNA in cells, but the ratio would be much larger if the host is a *Methanoperedens* species. The linkage between mini-Borgs and *Methanoperedens* is strongly supported by gene phylogenies (Figs. 2 and S5-9), so a genome copy number ratio of at least 600 : 1 is predicted for this case. Either way, the findings raise the possibility that the abundant mini-Borg DNA derives from a recent (or ongoing) viral-like bloom. Alternatively, the mini-Borgs may exist in the form of relict environmental DNA that is mostly protected by an as-yet-unknown mechanism. The strong dips in coverage near the termini of some mini-Borg genomes suggest that some of the DNA is old and unprotected. Borgs are not reported to display such patterns, perhaps because their genomes are protected by the much longer ITRs.

Notably, mini-Borgs encode some genes homologous to those involved in the *Methanoperedens* metabolism. One, replication factor C (RFC) small subunit, is encoded on the genomes of Saturn mini-Borgs (Fig. S5). RFC plays a critical role in DNA replication through loading the proliferating cell nuclear antigen that can tether DNA polymerase to the template thus promoting the processivity of DNA synthesis^40^. Archaeal RFC requires large and small subunits^41,42^. Without the large subunit, the mini-Borg small subunit could not be active^41^. By analogy with viruses/phages that have single subunits of multi-subunit complexes^15^, the RFC small subunits carried by mini-Borgs may boost the enzymatic activity of the host’s RFC.

Some genomes of mini-Borgs encode archaeal A-type ATP synthase subunit K with high similarity to *Methanopereden*s homologs (Fig. 2a). This subunit forms a ring-shaped structure embedded in the membrane and assists in proton translocation^43,44^. Given that mini-Borgs (and other ECEs) are unable to synthesize the entire ATPase complex, this subunit may facilitate the transportation of protons across the *Methanopereden*s cytoplasmic membrane to promote ATP production.

To our surprise, one *Methanoperedens* virus encodes two MHCs, which have never been detected in other virus/phage genomes, but they are ubiquitous and in multi-copy in Borg genomes^14^. Thus, MHCs in ECEs appear to be a more common phenomenon than previously reported. The virus-borne MHCs show high similarity to homologs in a *Candidatus* Methanoperedenaceae archaeon (Accession: GCA_014859785.1) that lacks the nitrate reductase complex^45^, suggesting these MHCs likely transfer electrons from menaquinone to poorly soluble Fe(III) and Mn(IV) oxyhydroxides^8,9^. Given the predicted extracellular localization of viral MHCs, and consistent with prior inferences for the significance of metabolic genes on phages^46,47^, we propose that viruses can augment electron transfer for *Methanoperedens* during viral production.

Some viruses of *Methanoperedens* carry a ThyA gene that is also encoded in the genomes of Borgs and plasmids (Fig. S23). In contrast, ANME-1 viruses encode non-homologous counterparts ThyX^48^. The difference can be explained by the metabolism of the hosts, as *Methanoperedens* use the ThyA pathway for thymidylate synthesis whereas ANME-1 mainly use the ThyX pathway^37^. The similarity of host- and virus-encoded genes is likely due to gene exchange between hosts and their associated ECEs. The findings for the thymidylate genes further extend the observation that the genes of viruses tend to function in the context of host-encoded pathways. Although true also of some Borg-encoded genes, Borgs differ in that their genomes also encode some functional complexes and pathways, including metabolic genes not encoded in their host’s genomes^12^.

Some of the ECEs we described could not be classified. Certain of these circular extrachromosomal elements encode TnpBs (Fig. 4), proteins of great interest for the evolution of CRISPR-Cas systems^49^. TnpB in bacteria represents the minimal structural and functional core of Cas12 endonucleases that cleaves DNA strands and thus is used to edit genomes in human cells^50^. Unlike Cas12s, which are guided by spacer-derived RNA, TnpB is normally directed by the RNA transcribed from the right end of the transposon^51,52^. Reprogramming for genome editing consequently requires a modification of the TnpB transposon instead of designing a simple RNA guide. However, the ECE TnpB is encoded adjacent to CRISPR arrays and one locus is highly diversified at the population level, implying that it is actively integrating spacers (Fig. 4). This raises the possibility that this TnpB may be CRISPR spacer-directed. A number of genetic elements appear to be targeted by the spacers, but a target was not identified for the right end of the TnpB transposon, supporting the hypothesis that the ECE TnpB is directed by CRISPR spacers. If proven biochemically, genome editing tools developed using the ECE system may be of utility, especially as this TnpB is smaller than Cas proteins and guide RNA is easier to design compared to modifying transposons.

Given the coexistence of *Methanoperedens* and multiple ECE types, there are many opportunities for lateral gene transfer. We observed evidence for the movement of at least six genes between *Methanoperedens* and two distinct ECEs, and five genes between *Methanoperedens* and more than two ECEs, including transposases shared among *Methanoperedens* and all of its associated ECEs (Fig. 5). Thus, there appears to exist a complex gene exchange network in which *Methanoperedens* serves as the hub (Fig. 5).

**Fig. 5.**
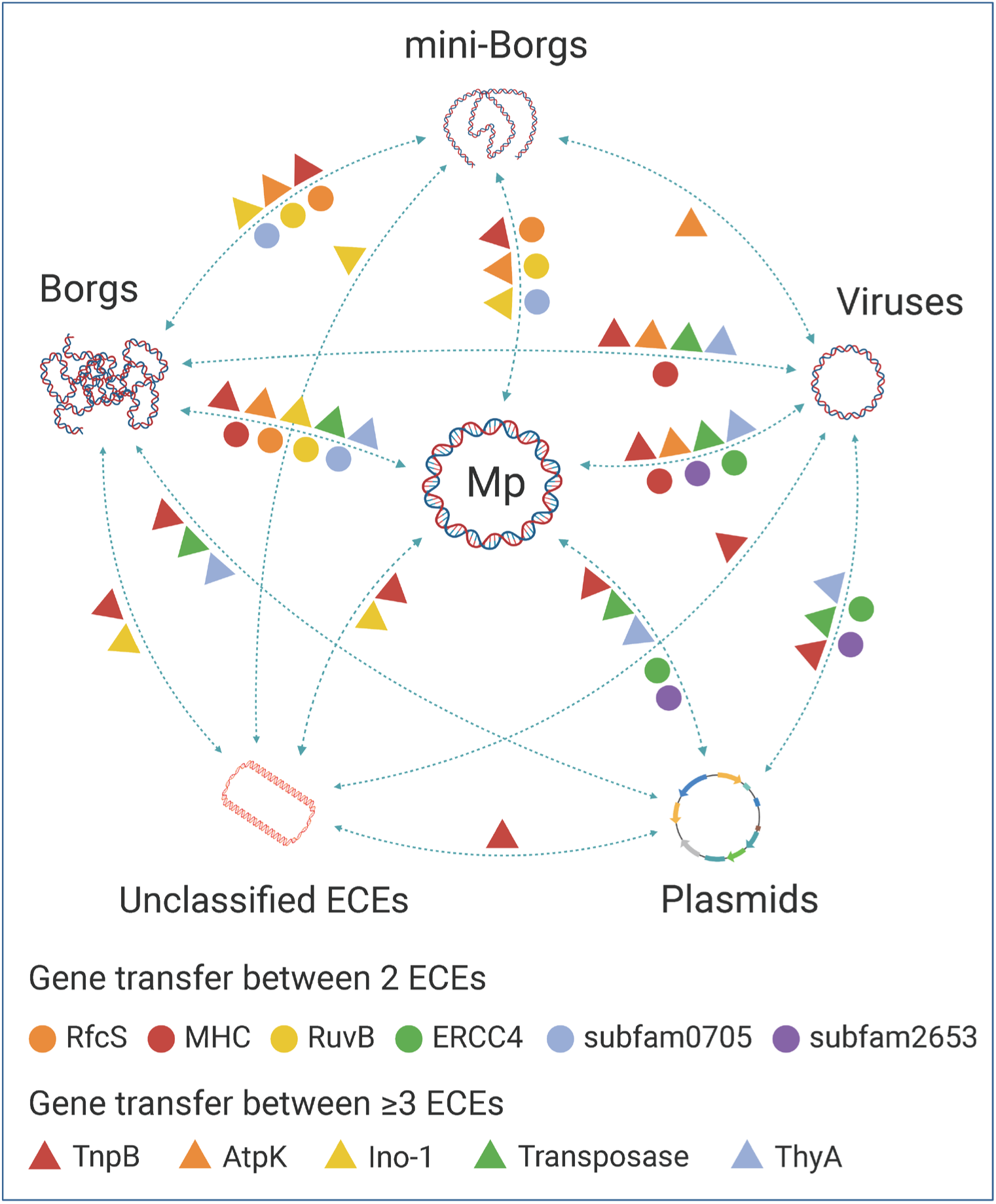
Evidence for lateral gene transfer amongst *Methanoperedens* and associated ECEs based on gene phylogenies. Colored circles and triangles indicate genes for which closely related homologs were identified in different genomes. Abbreviations provide examples of transferred genes. Detailed phylogenies for each gene can be found in the main and supplementary figures.

In conclusion, we greatly expand the repertoire of ECEs that our data strongly suggest are associated with *Methanoperedens.* The extent to which these ECEs impact *Methanoperedens* activity, especially the rate at which it oxidizes methane, remains unknown. It is clear that their gene contents have the potential to augment those of their hosts, and in multiple functional categories. However, if our inference of virus-like dynamics is correct, they may also decimate *Methanoperedens* populations and thus reduce the feedback to methane emissions. In combination with prior work on Borgs, the results indicate that the ECEs of *Methanoperedens* are highly variable in their abundances, genome sizes, genome architectures, and gene contents.

## Methods

### Identification and genome curation of *Methanoperedens*-associated ECEs

Metagenomic datasets on ggKbase (ggkbase.berkeley.edu) and the latest IMG/VR v4 database were used for searching candidates of novel ECEs potentially associated with *Methanoperedens* first based on taxonomic profiles and GC contents. Recruited contigs were manually curated to extend and remove assembly errors using Geneious Prime 2022.2.2 (https://www.geneious.com), as detailed in our previous paper^53^. Replichores of completed ECEs were predicted according to the GC skew and cumulative GC skew calculated by the iRep package (gc_skew.py)^18^. For circular ECEs, the origin of replication was moved to the start of genomes. Genome relatedness between the host *Methanoperedens* and associated ECEs was compared by global proteome-based alignment and visualized in a BIONJ tree (https://www.genome.jp/digalign/). Identified ECEs with viral structural genes were further designated as viruses.

### Correlation analyses for mini-Borgs

Abundances of each *Methanoperedens* and mini-Borg genome was calculated across 68 wetland samples using CoverM v0.6.1 (https://github.com/wwood/CoverM). For mini-Borgs, the “contig” mode was run with minimum reads identity of 95% and minimum aligned percent of 75%. For *Methanoperedens*, genomes were first dereplicated at average nucleotide identity of 95% using dRep v3.4.0^54^, and then calculated using the “genome” mode with the same cutoff parameters as above. Correlations were calculated using Spearman’s rank-order correlation metric with the “Hmisc” package in R^55,56^.

### Classification and comparison of *Methanoperedens*-associated viruses

The host of identified viruses was predicted by: 1) CRISPR spacer matches, in which CRISPR-Cas loci of *Methanoperedens* were predicted using CRISPRCasTyper v1.8.0^57^ and recruited spacers were matched against the viruses with a minimum similarity of 95% using BLAST^58^; 2) Blast-based comparisons, where viruses were aligned with microbial genomes at a maximum e-value 1×10^−3^, minimum identity 80%, and minimum alignment length 500 nt; 3) and other virus-specific host prediction tools including VirHostMatcher (VHM)^59^, WIsH^60^, Prokaryotic virus Host Predictor (PHP)^61^, and RaFAH^62^, which are all integrated in the iPHoP v1.2 that generates an iPHoP-RF value simultaneously^63^. The first two prediction methods were also applied to other associated ECEs besides viruses.

Newly identified *Methanoperedens* viruses were compared to available prokaryotic viruses through: 1) gene-sharing networks^64^ with viral references including RefSeq viral genomes (release 217), Asgard and ANME-1 archaeal viruses^48,65^; and 2) comprehensive blast against the latest IMG/VR v4 database^23^. Phylogenetic classification was predicted based on genome-wide similarities using ViPTree^66^. By comparing the genetic distances between and across halovirus families, we assigned *Methanoperedens* viruses into eight different families and designated them asteroid names according to genome sizes. Gene cluster and alignment among *Methanoperedens* viruses were analyzed using Clinker v0.0.27^67^, after the generation of genbank files by Prokka v1.14.6^68^.

### Prediction, annotation, and phylogenetic analyses of genes inside *Methanoperedens*-associated ECEs

Genes in *Methanoperedens*-associated ECEs were first predicted using Prodigal v2.6.3^69^, and then searched against 1) KEGG^21^, UniRef100^20^, and UniProt^20^ databases by USEARCH v10^70^, and 2) NCBI nr database by BLAST^58^. Functional domains inside the translated protein sequences were scanned against InterPro’s member databases using InterProScan v5^71,72^. Subcellular localization was predicted by PSORT v2.0 using archaeal mode^73^. Proteins of interest were aligned with close references using MAFFT v7.453^74^, followed by an automatic trimming with trimAl v1.4.rev15^75^. Sequence alignments were further used to construct phylogenetic trees using IQ-TREE v1.6.12 with best-fit models determined automatically^76^. Generated trees were decorated on the iTOL webserver^77^.

### Structural predictions and comparisons of TnpB

The TnpB sequence from the representative unclassified ECE was structurally modeled using AlphaFold2 via LocalColabFold with default parameters^78^. Structural homologs of the predicted structure were identified using Foldseek^79^. Finally, the TnpB and recruited homologous structures were superimposed, compared, and visualized in PyMOL (v2.3.4)^80^.

## Data availability

Metagenomic sequencing reads related to *Methanoperedens* ECEs are available under NCBI BioProject: PRJNA999944. Prior to publication, the genomes described in this study can be accessed at: https://ggkbase.berkeley.edu/project_groups/methanoperedens_ece. Please note that it is necessary to sign up as a user (simply provide an email address) in order to download the data.

## Acknowledgments

The authors would like to thank Jennifer A. Doudna and Ben Adler for helpful discussion of the TnpB-associated CRISPR complex. Funding for this research was provided by the Bill and Melinda Gates Foundation (Grant Number: INV-037174 to J.F.B). The findings and conclusions are those of the authors and do not necessarily reflect positions or policies of the Bill and Melinda Gates Foundation. Funding was also provided via the Innovative Genomics Institute Climate fund philanthropic donation to JFB, a DFG postdoctoral fellowship to M.C.S. (Project Number: 447383558 to M.C.S.), and The Ministry of Economy, Trade and Industry of Japan as “The project for validating near-field system assessment methodology in geological disposal system” (2022 FY, Grant Number: JPJ007597).

## Author contributions

The study was designed by L-D.S. and J.F.B. Binning and genome curation and analysis were performed by L-D.S. and J.F.B. Proteome, and phylogenetic analyses were carried out by L-D.S. Correlation analyses were performed by L-D.S. J.W-R. provided the Corona Mine sequences and computational support. M.C.S. contributed to proteome analyses and P.I.P. carried out the protein structural analyses. LX.C contributed to TnpB and CRISPR array analyses. Y.A. provided the Horonobe metagenomic sequences and S.L. and R.S. contributed to the data handling and bioinformatic analyses. L-D.S. and J.F.B. wrote the manuscript with input from all the authors.

## Supplementary Information

**Fig. S1.**
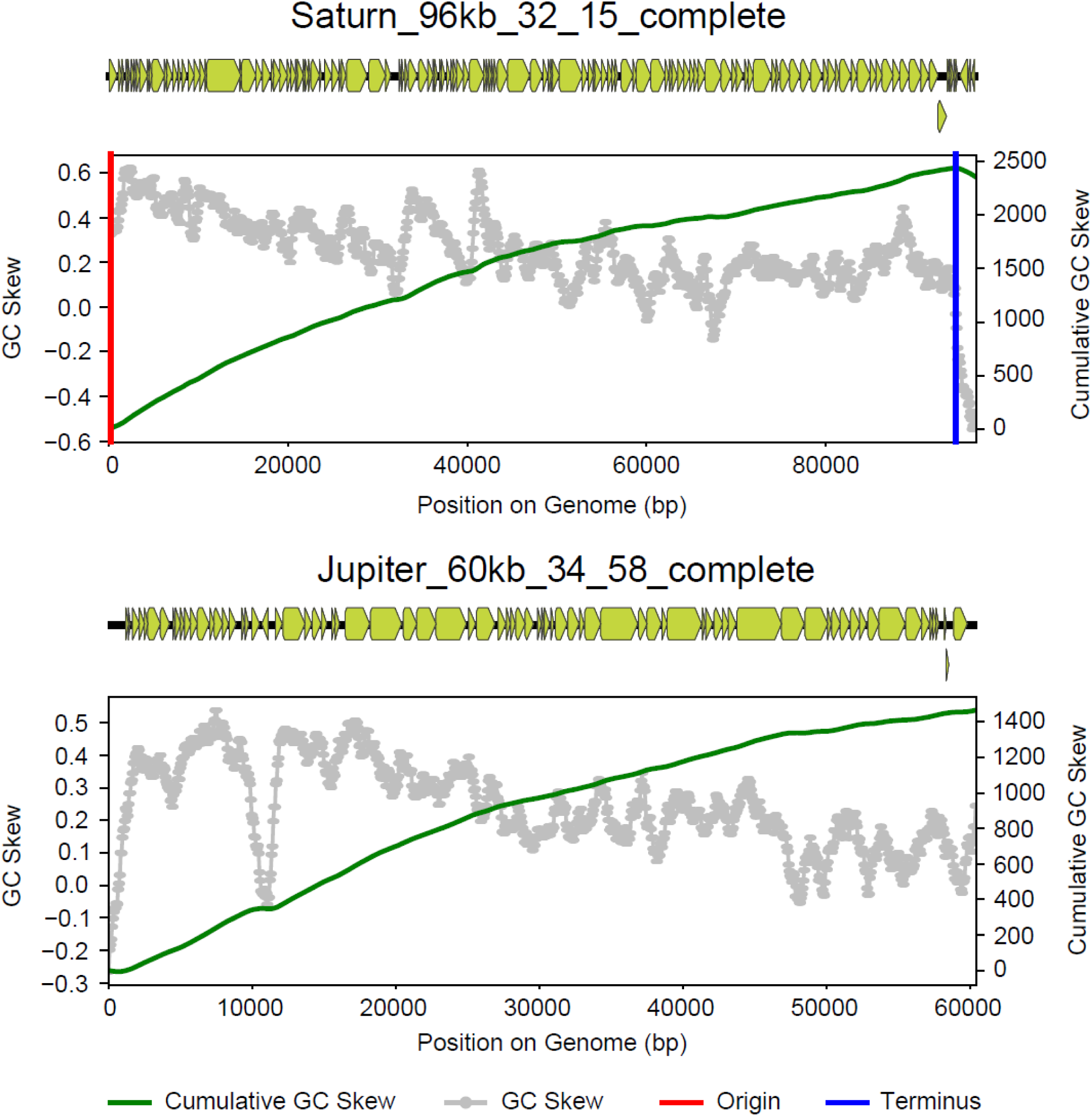
Genomic architectures and predicted replichores of representative complete mini-Borgs. Yellow blocks indicate predicted genes while red and blue lines indicate predicted replication origin and terminus, respectively. Gray dots and green lines show GC skew and cumulative GC skew across genomes.

**Fig. S2.**
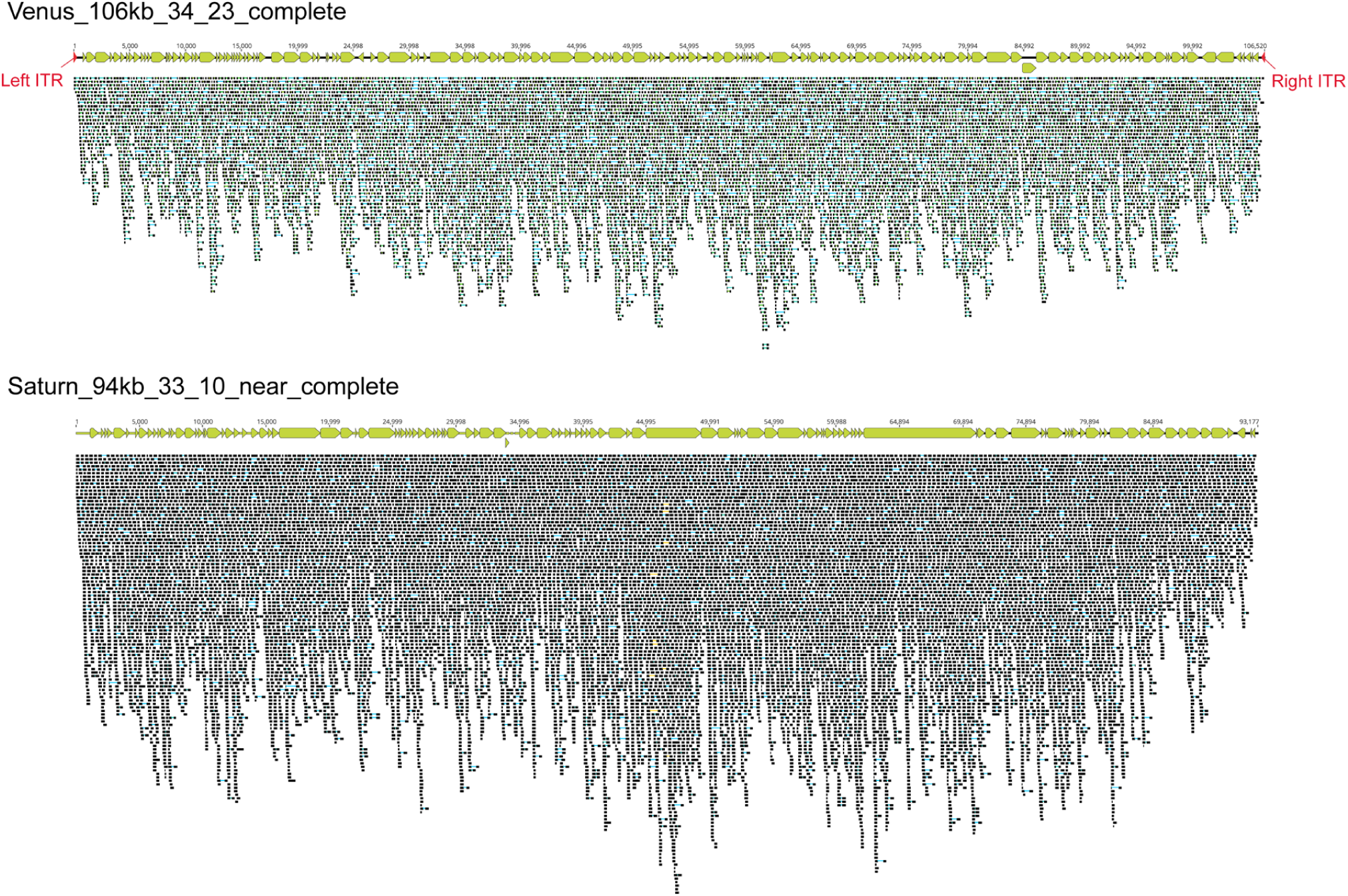
Coverage profile of representative complete and near-complete mini-Borgs. Yellow blocks indicate predicted genes. Red arrows in the Venus_106kb_34_23_complete denote inverted terminal repeats (ITR). Mapped reads are depicted below the genomes.

**Fig. S3.**
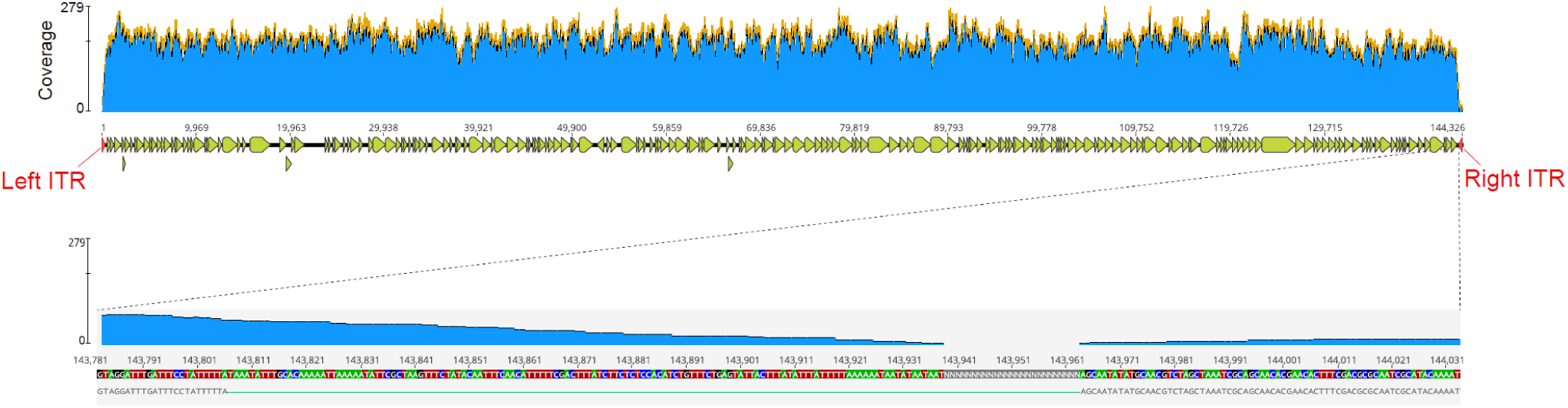
Genome architecture of Jupiter_144kb_32_101_near_complete supported by paired reads. Yellow blocks and red arrows indicate predicted genes and ITR. Blue area denotes the genome coverage. The bottom inset only shows the paired reads spanning the gap region, which strongly supports the presence of the inverted terminal repeat.

**Fig. S4.**
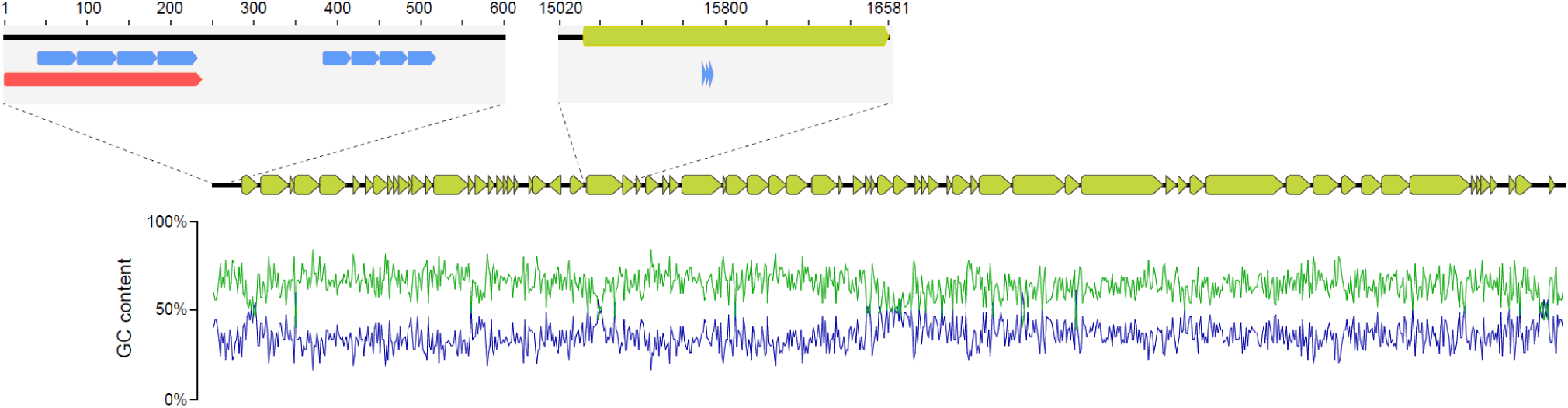
Examples of tandem repeats in the Jupiter_54kb_36_306_complete genome. Yellow blocks indicate predicted genes; red and blue blocks denote, respectively, left inverted terminal repeat and perfect tandem repeats distributed in various regions.

**Fig. S5.**
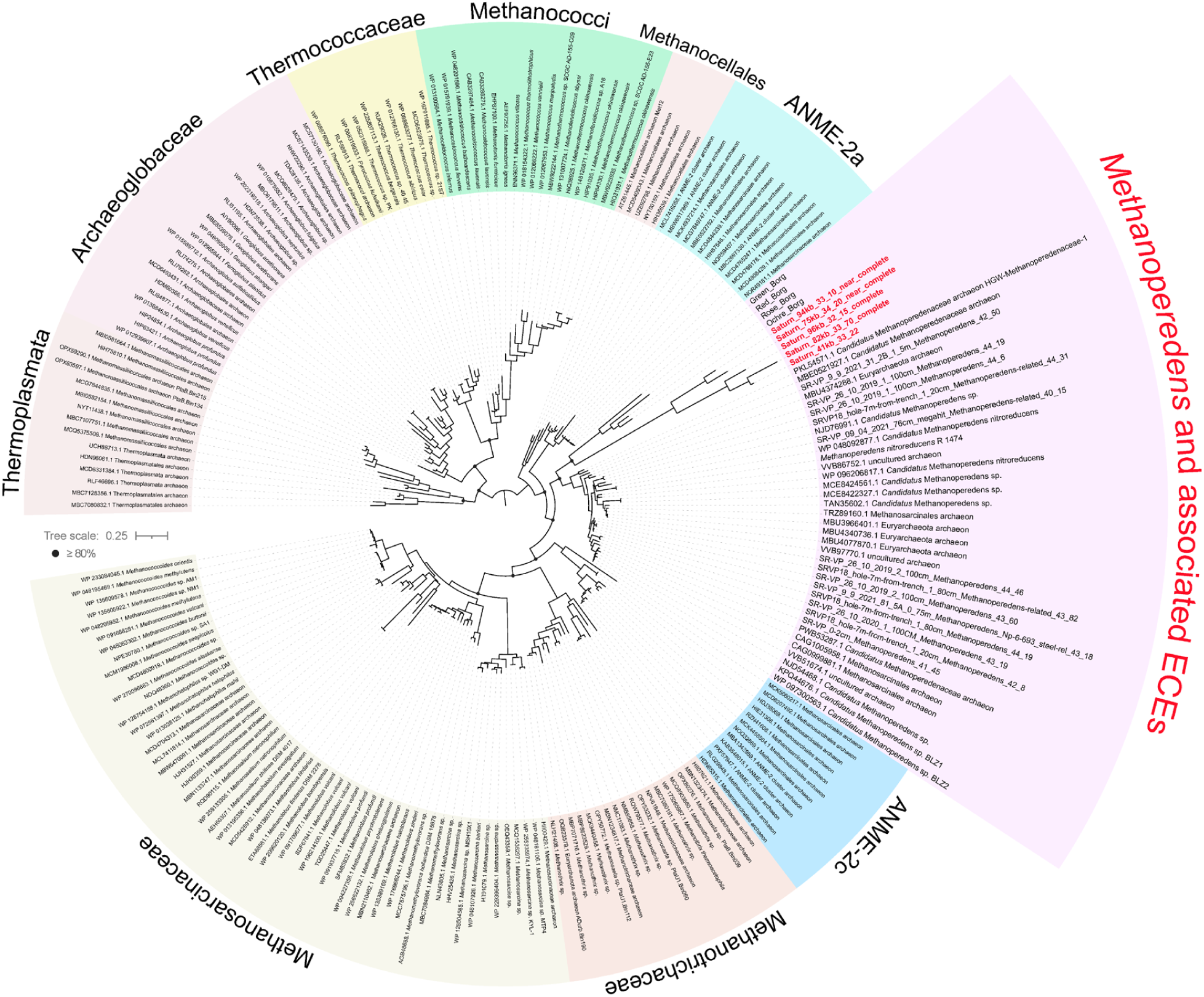
Phylogeny of replication factor C small subunit (RfcS) in *Methanoperedens* and associated ECEs. Homologous references were recruited from the NCBI nr database by blast. Arc colors indicate different taxonomic clades. Proteins found in mini-Borgs are highlighted in red. The tree was mid-point rerooted and support values were calculated based on 1000 replicates.

**Fig. S6.**
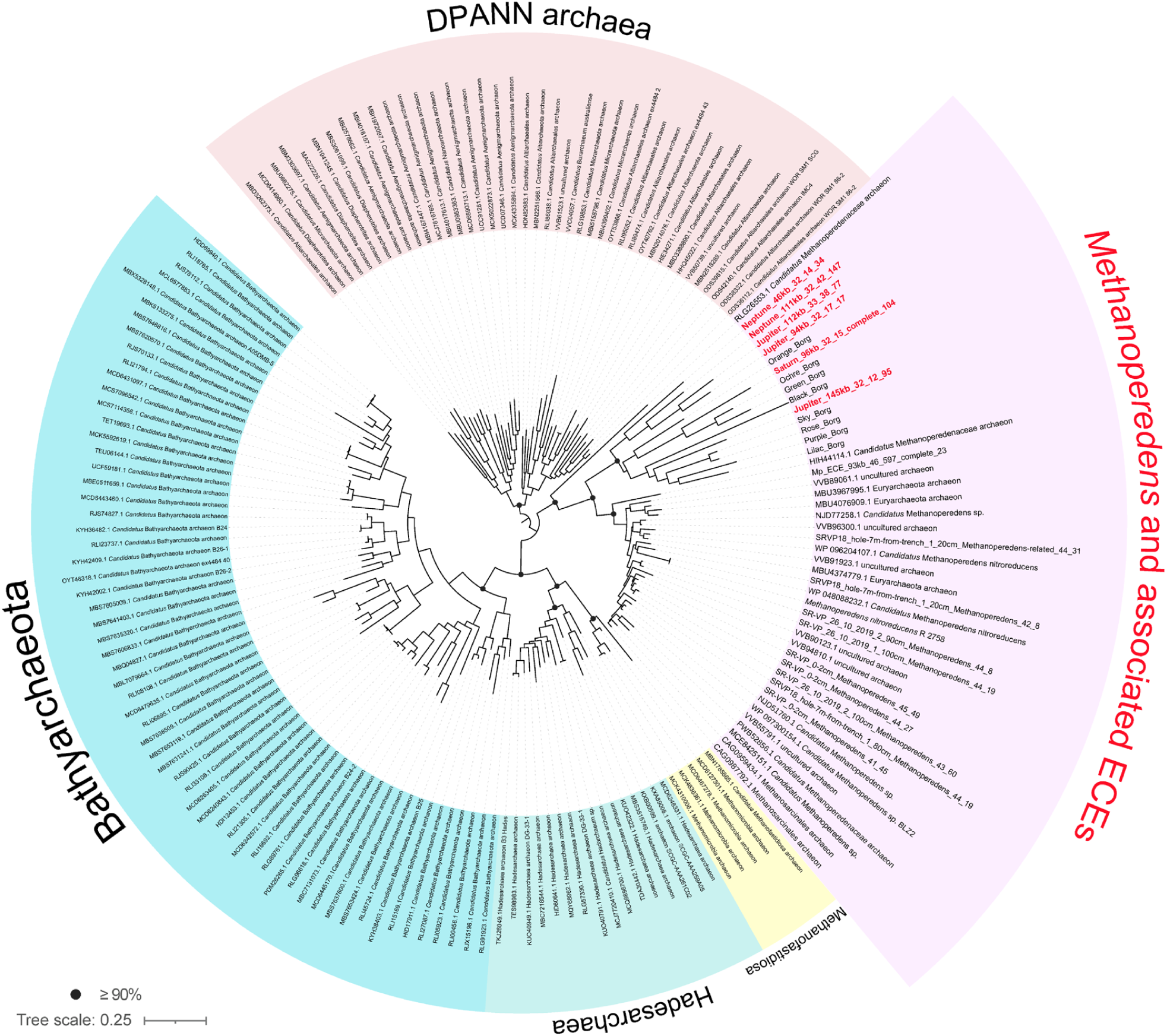
Phylogeny of myo-inositol-1-phosphate synthase (Ino-1) in *Methanoperedens* and associated ECEs. Homologous references were recruited from the NCBI nr database. Arc colors indicate different taxonomic clades. Proteins found in mini-Borgs are highlighted in red. The tree was mid-point rerooted and support values were calculated based on 1000 replicates.

**Fig. S7.**
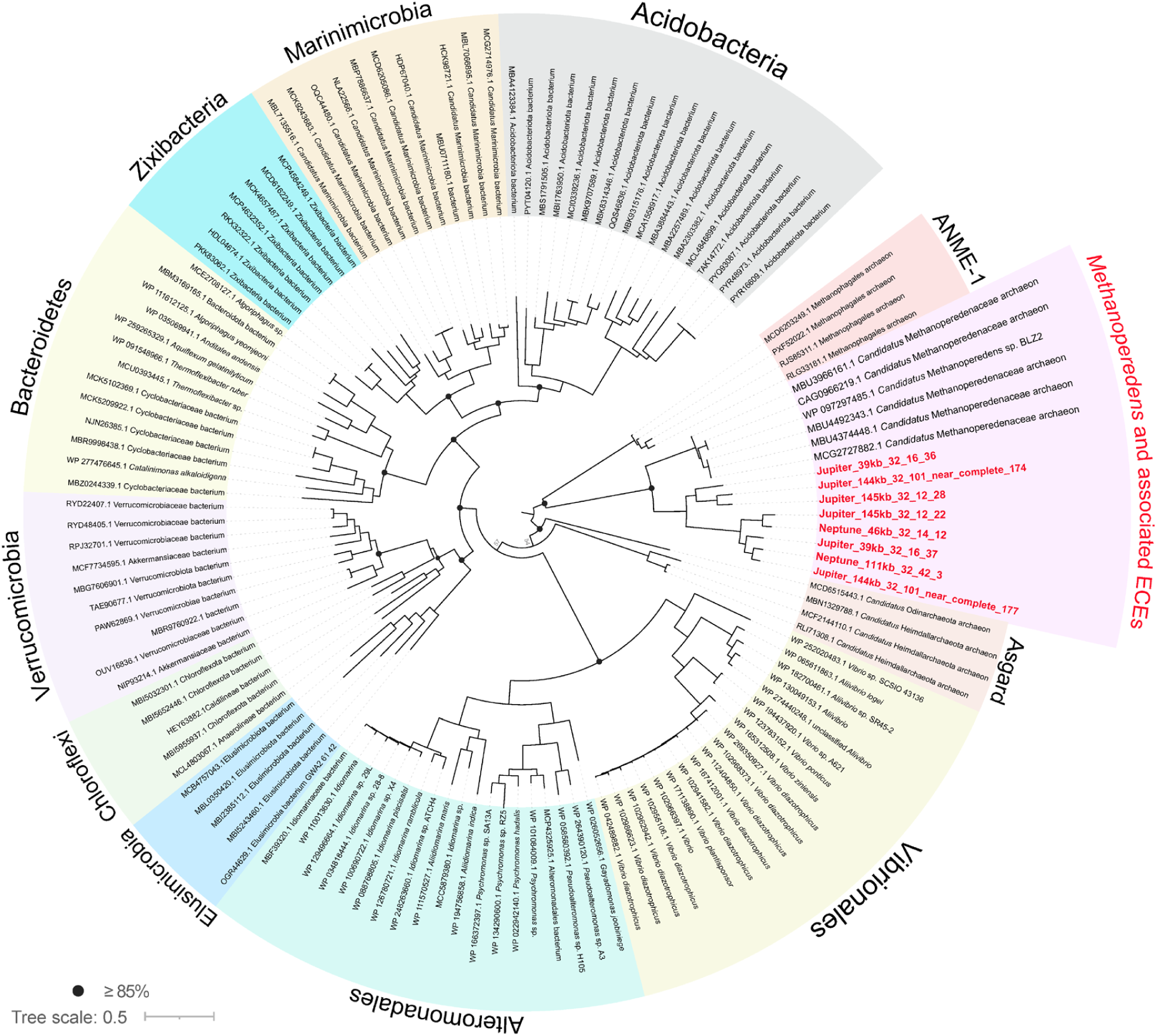
Phylogeny of MBL (Metallo-β-lactamase) fold metallo-hydrolase in *Methanoperedens* and associated ECEs. Homologous references were recruited from the NCBI nr database. Arc colors indicate different taxonomic clades. Proteins found in mini-Borgs are highlighted in red. The tree was mid-point rerooted and support values were calculated based on 1000 replicates.

**Fig. S8.**
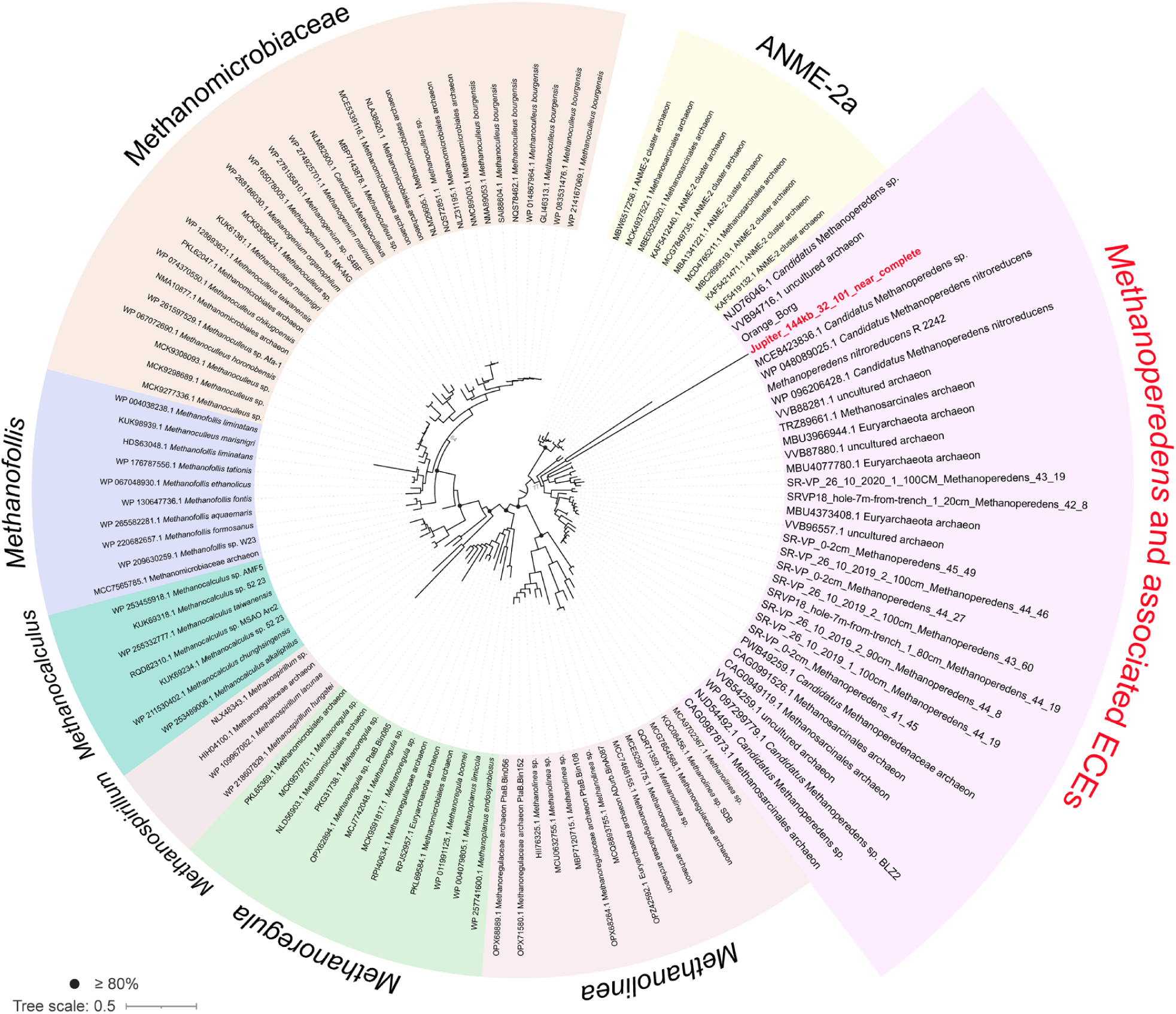
Phylogeny of DNA helicase RuvB in *Methanoperedens* and associated ECEs. Homologous references were recruited from the NCBI nr database. Arc colors indicate different taxonomic clades. Proteins found in mini-Borgs are highlighted in red. The tree was mid-point rerooted and support values were calculated based on 1000 replicates.

**Fig. S9.**
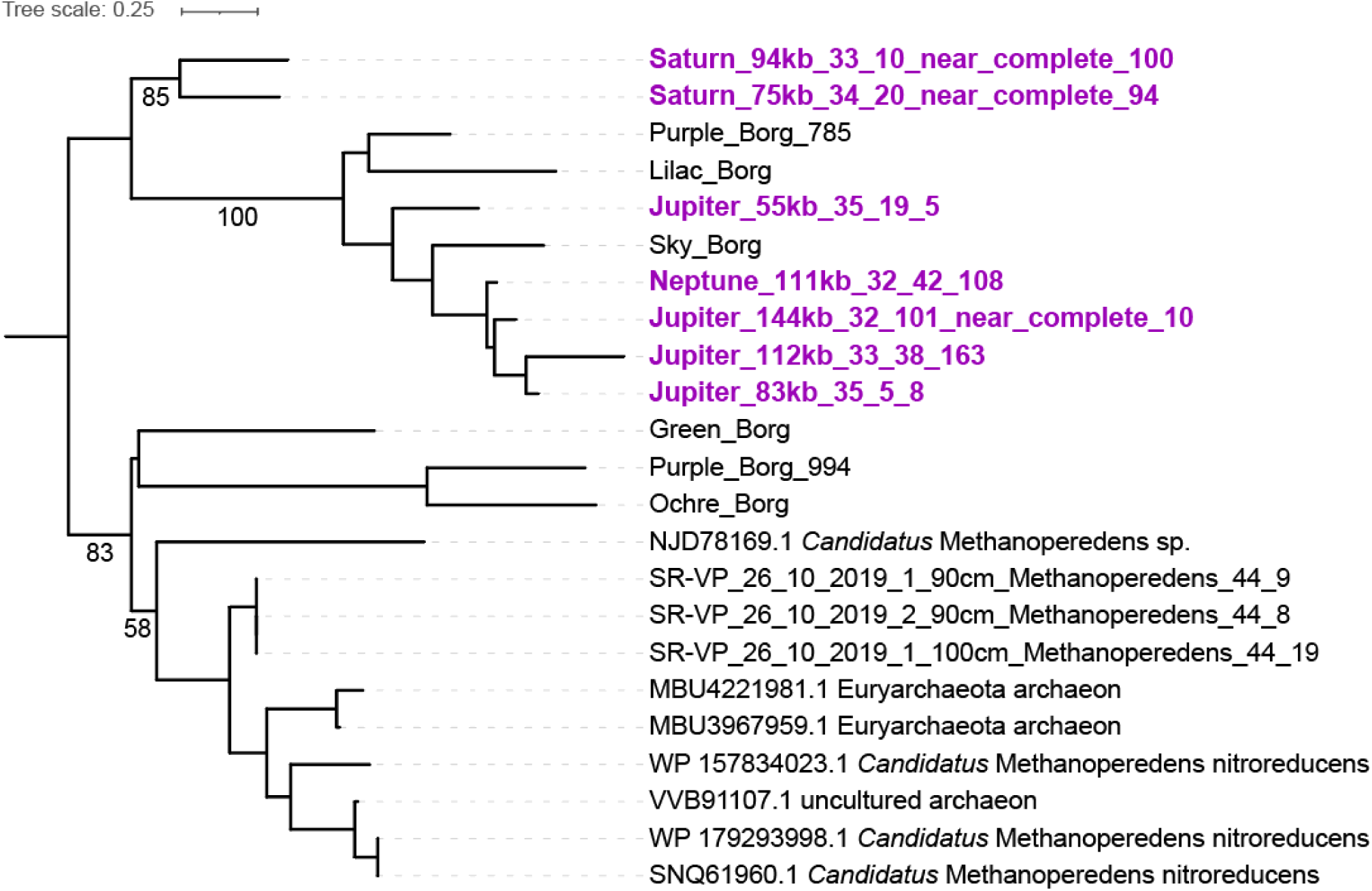
Phylogeny of a hypothetical protein (subfam0705) in *Methanoperedens* and associated ECEs. The only related sequences from the NCBI nr database are from *Methanoperedens*. Proteins found in mini-Borgs are highlighted in purple. The tree was mid-point rerooted and support values were calculated based on 1000 replicates.

**Fig. S10.**
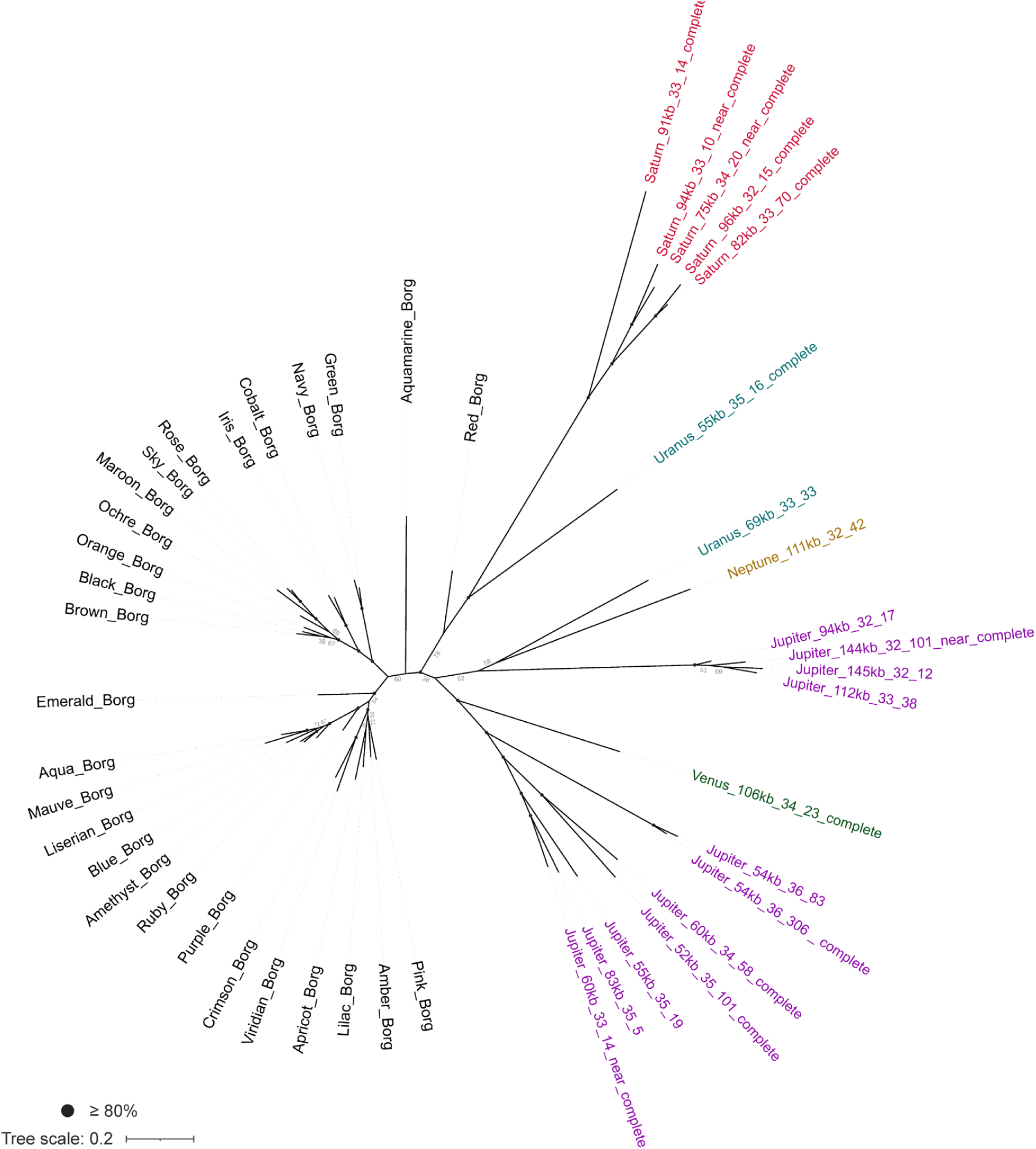
Phylogeny of Borgs and mini-Borgs inferred from a concatenation of two marker proteins. Blastp of markers against the NCBI database recruited no significant hits, therefore no other sequences are included in the tree. Support values were calculated based on 1000 replicates.

**Fig. S11.**
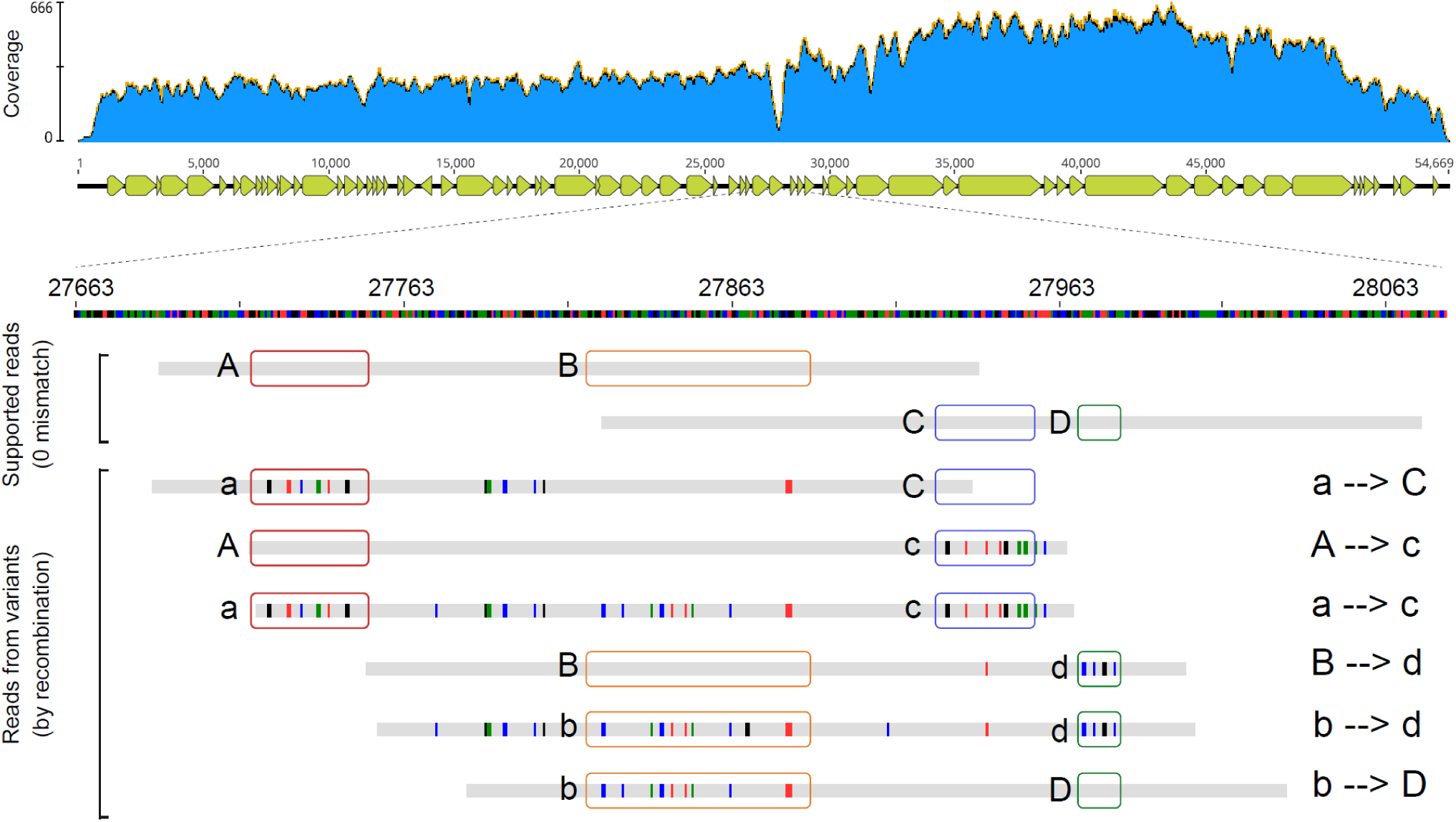
An example of recombination in the Jupiter_54kb_36_306_complete genome. The obvious dip in the coverage profile (blue area in the upper panel) suggests a region for which the more divergent reads were not recruited. Sequence blocks surrounded by colored rectangles indicate alleles that are linked in a variety of configurations, consistent with recombination involving mini-Borg variants.

**Fig. S12.**
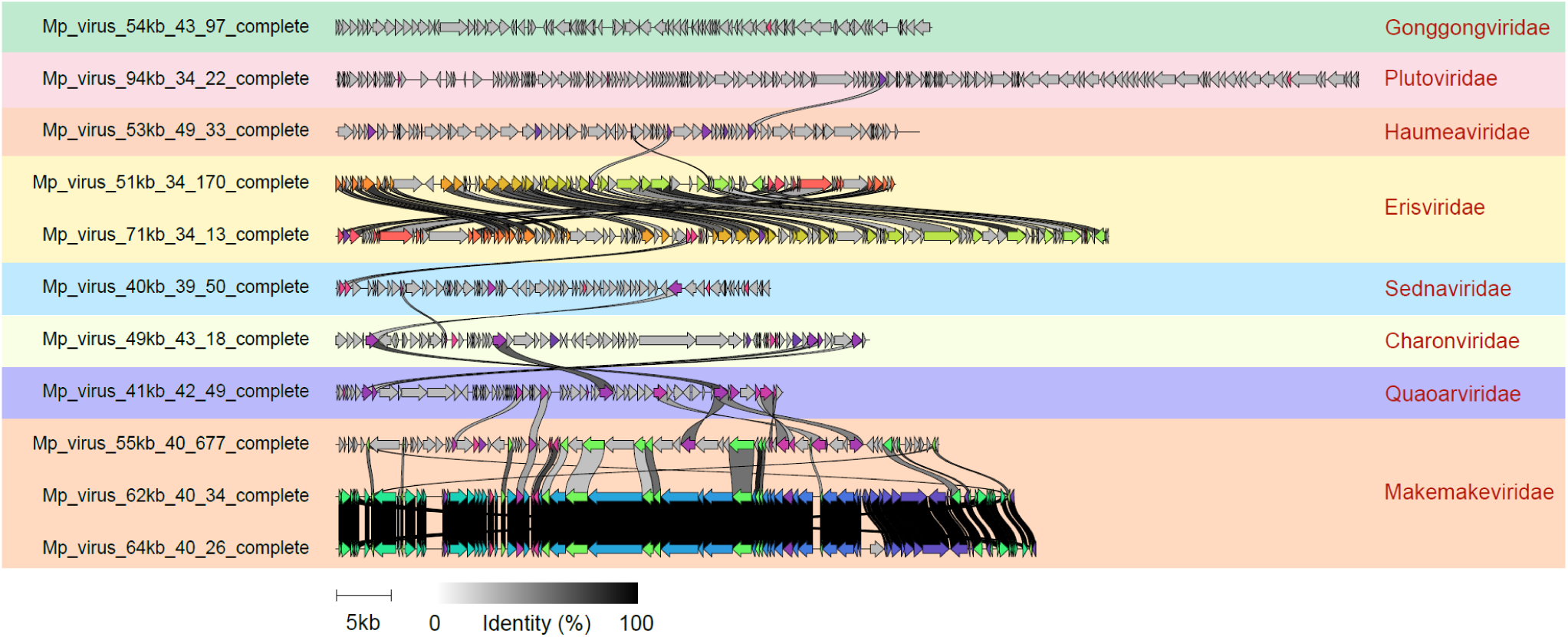
Genome alignment of 11 circularized *Methanoperedens* viruses belonging to eight families. Colored shadows indicate different virus families. The shade of gray lines that connect genes between genomes indicates pairwise amino acid identity. Gray arrows indicate proteins without homologs in other viruses.

**Fig. S13.**
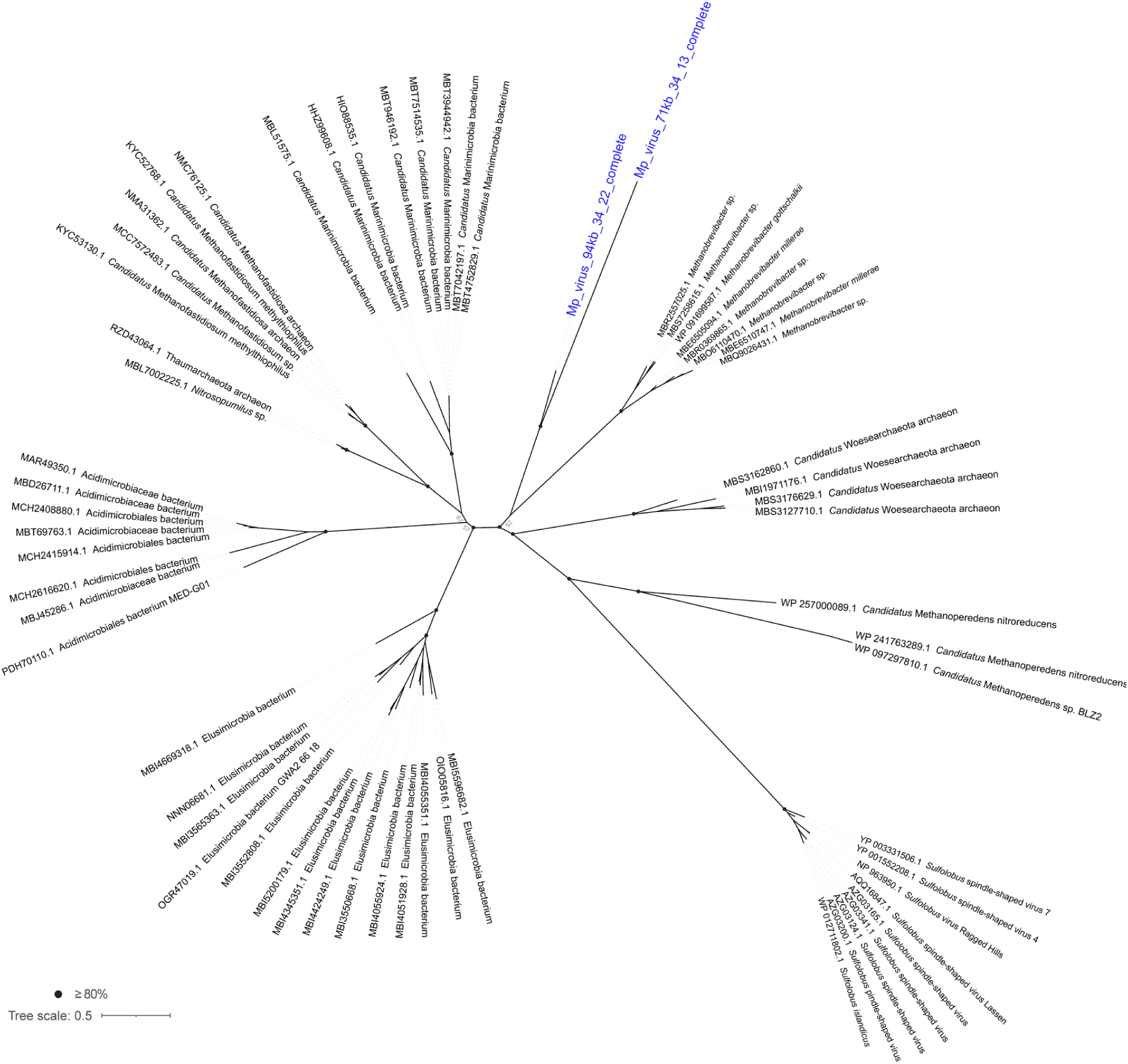
Phylogenetic tree of Cas4 proteins. Proteins in blue are from *Methanoperedens* viruses. Homologs were recruited from the NCBI nr database. Support values were calculated based on 1000 replicates.

**Fig. S14.**
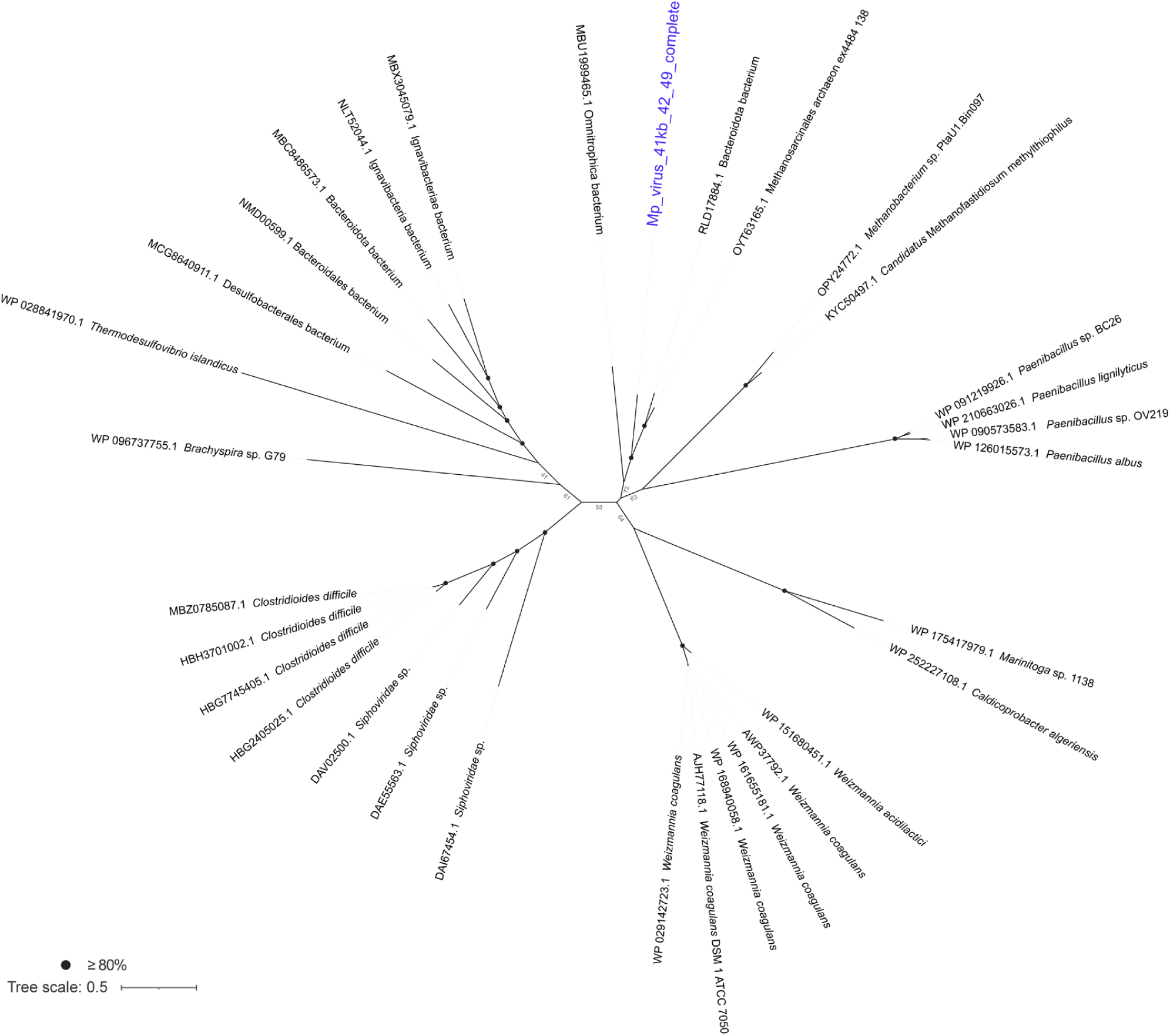
Phylogenetic tree of Type IV-B Csf1 (Cas8) proteins. The protein in blue is from a *Methanoperedens* virus. Homologs were recruited from the NCBI nr database. Support values were calculated based on 1000 replicates.

**Fig. S15.**
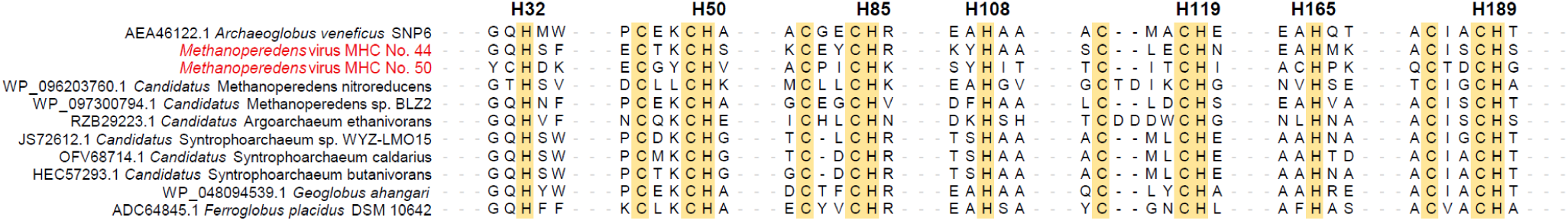
Sequence motifs conserved in MHC homologs forming extracellular cytochrome nanowires. Function and structure of the reference at the top are experimentally resolved. Coordinates indicate the residue position in the referenced protein belonging to *Archaeoglobus veneficus*.

**Fig. S16.**
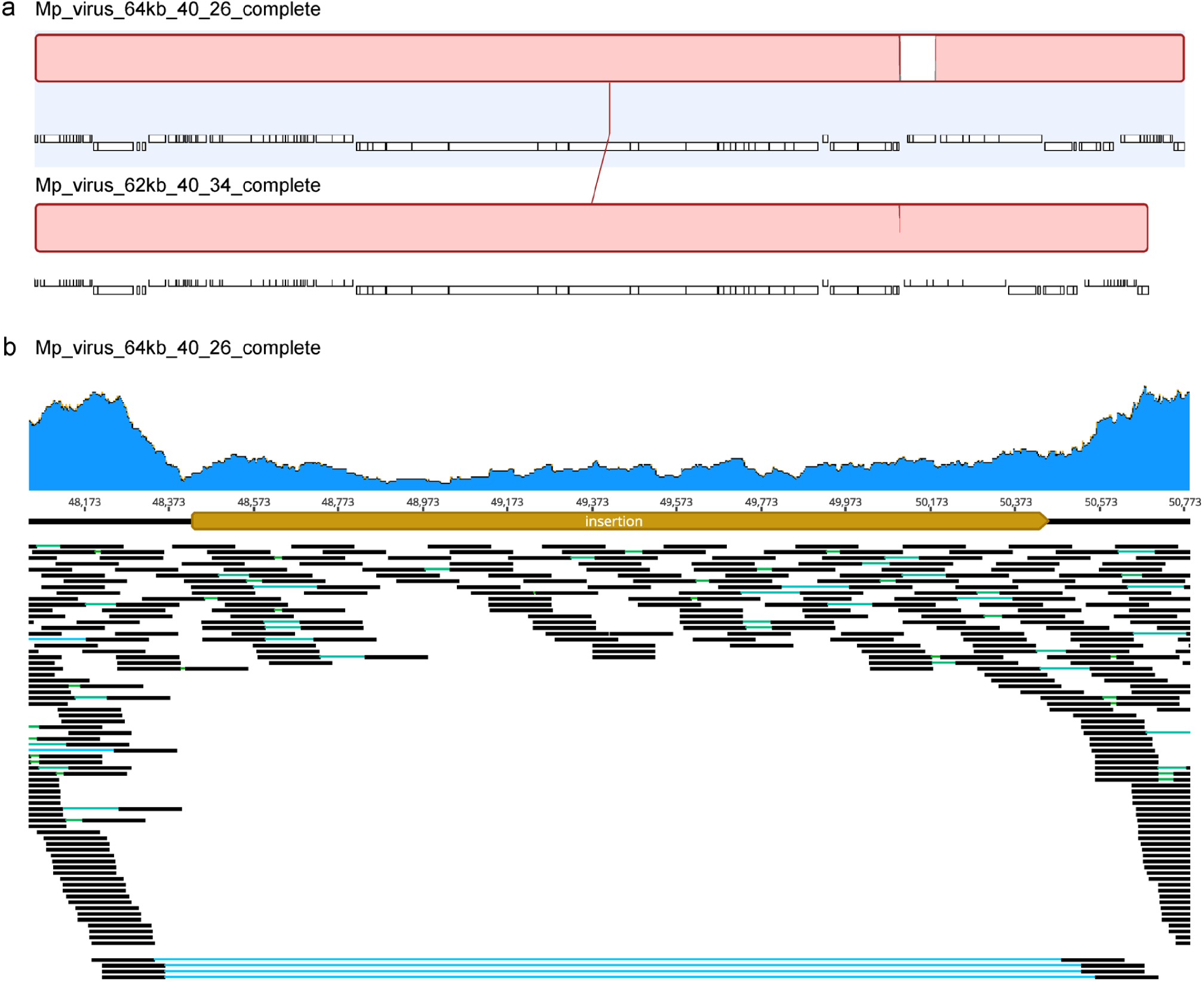
A ∼2 kb region integrated into the *Methanoperedens* viruses. **A.** Whole genome alignment of two closely related *Methanoperedens* viruses. The gap in the top genome corresponds with an insertion to the lower sequence. **B.** Read mapping shows paired reads spanning the inserted region.

**Fig. S17.**
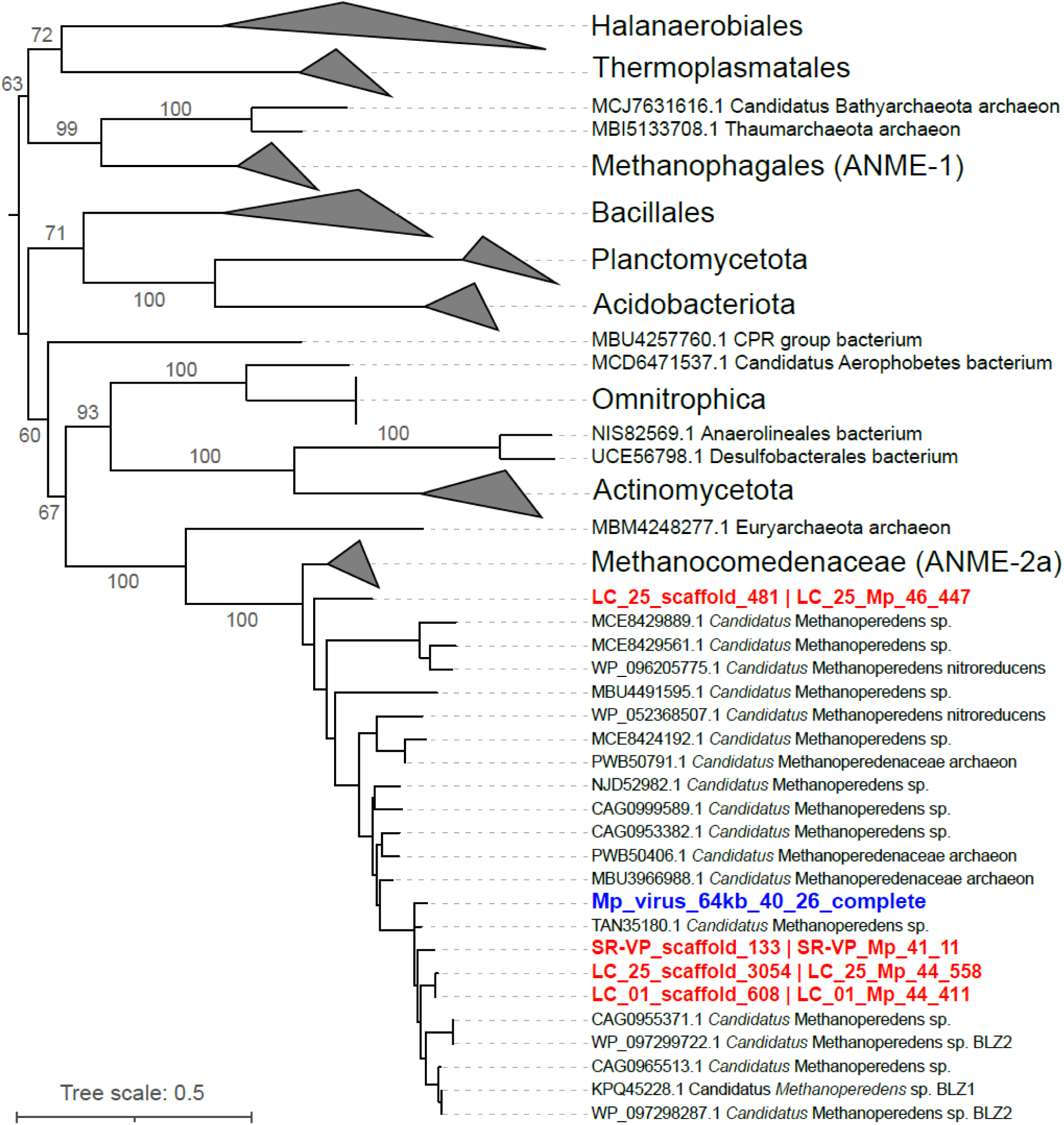
Phylogenetic tree of group II intron reverse transcriptase. Red names indicate proteins from *Methanoperedens* genomes (see Fig. S18) and blue names indicate proteins from viruses. The tree was mid-point rerooted and support values were calculated based on 1000 replicates. Similarity between *Methanoperedens* and virus sequences supports the inference that the viruses replicate in *Methanoperedens*.

**Fig. S18.**
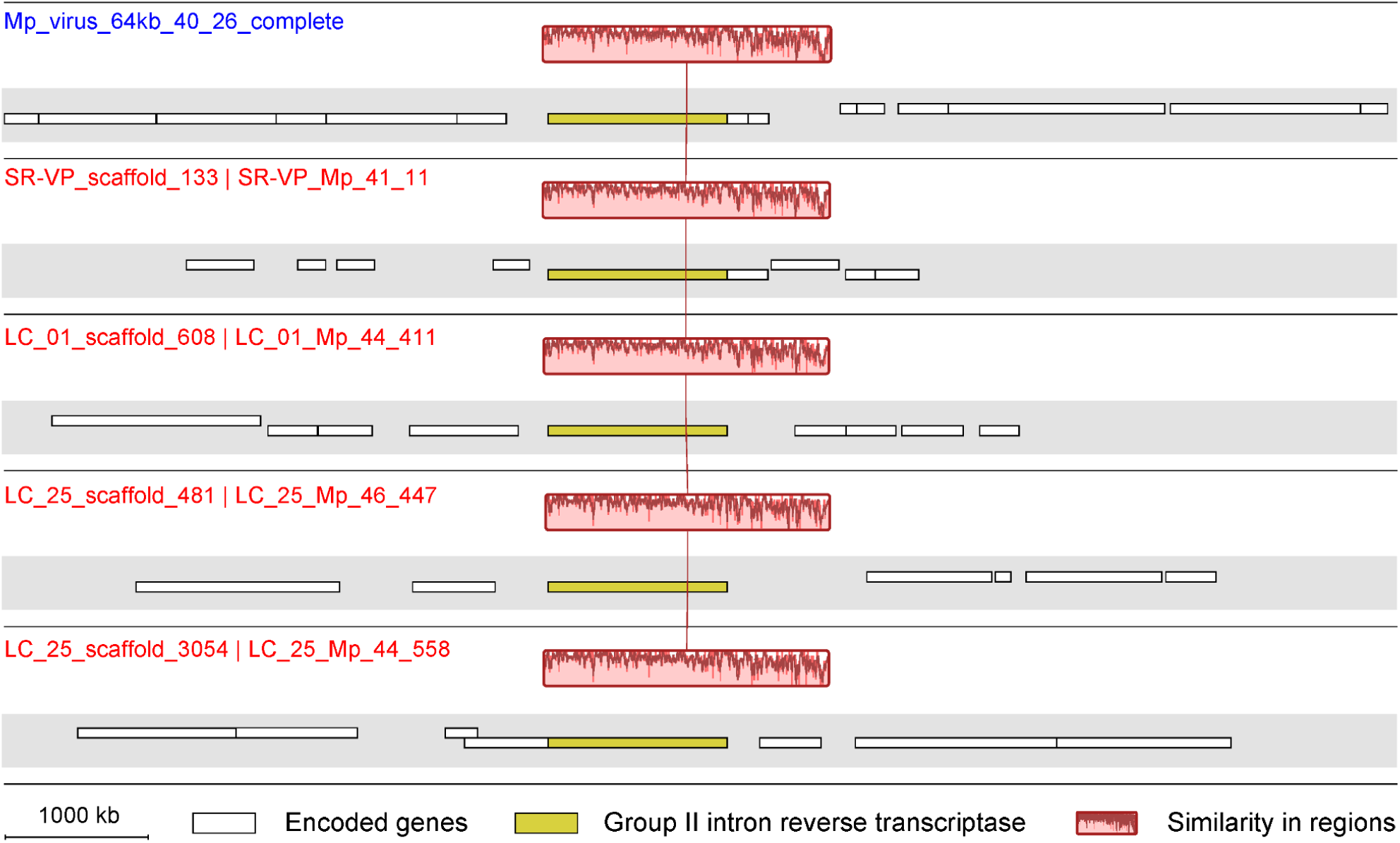
Genomic sequence comparison of intron-located regions. The virus sequence is shown at the top (blue name) and *Methanoperedens* genomes below (red names).

**Fig. S19.**
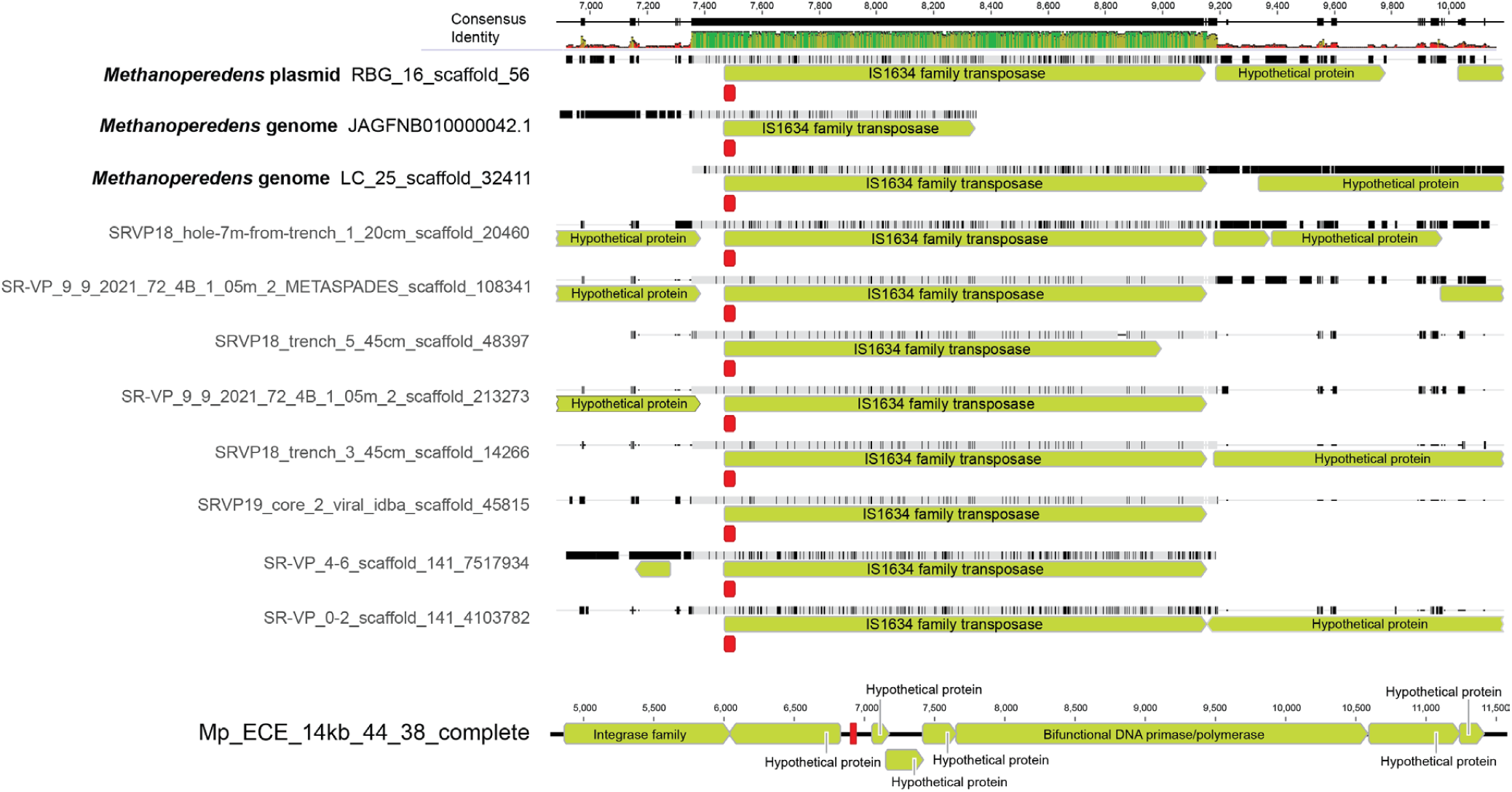
Regions that are targeted by TnpB-associated CRISPR spacers of Mp_ECE_93kb_46_597_complete. Protospacer regions corresponding to spacers are shown in red. “RBG_16_scaffold_56” is from a *Methanoperedens* plasmid (CP113844). “JAGFNB010000042.1” and “LC_25_scaffold_32411” is from *Methanoperedens* genomes, and others are unbinned. The bottom row is a *Methanoperedens*-associated, circularized, but unclassified ECE.

**Fig. S20.**
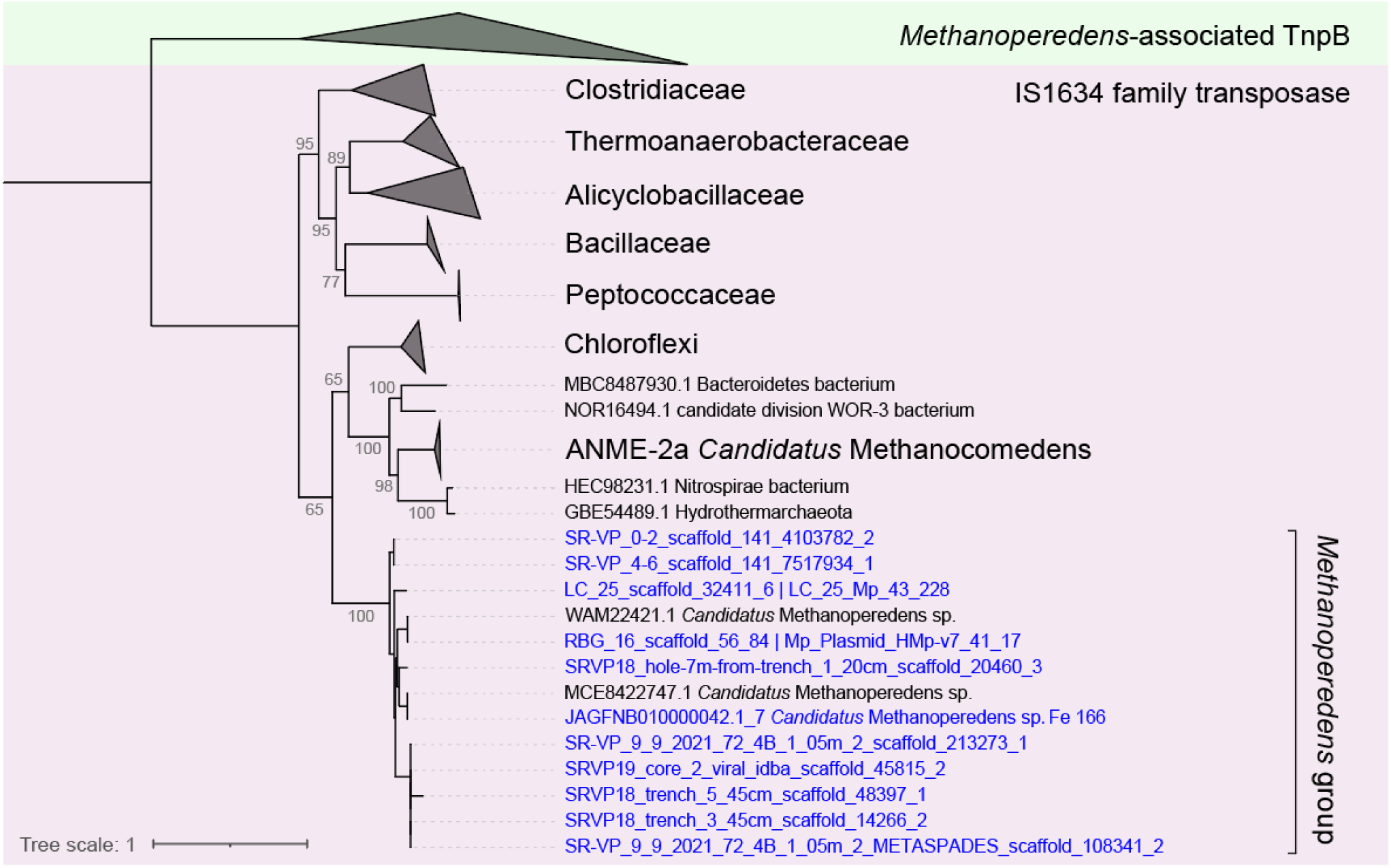
Comparison between transposon-associated TnpB and IS1634 family transposases. Transposases targeted by CRISPR arrays of the *Methanoperedens* ECE are marked in blue. Details of the TnpB group can be found in Fig. S21 below. The tree was mid-point rerooted and support values were calculated based on 1000 replicates.

**Fig. S21.**
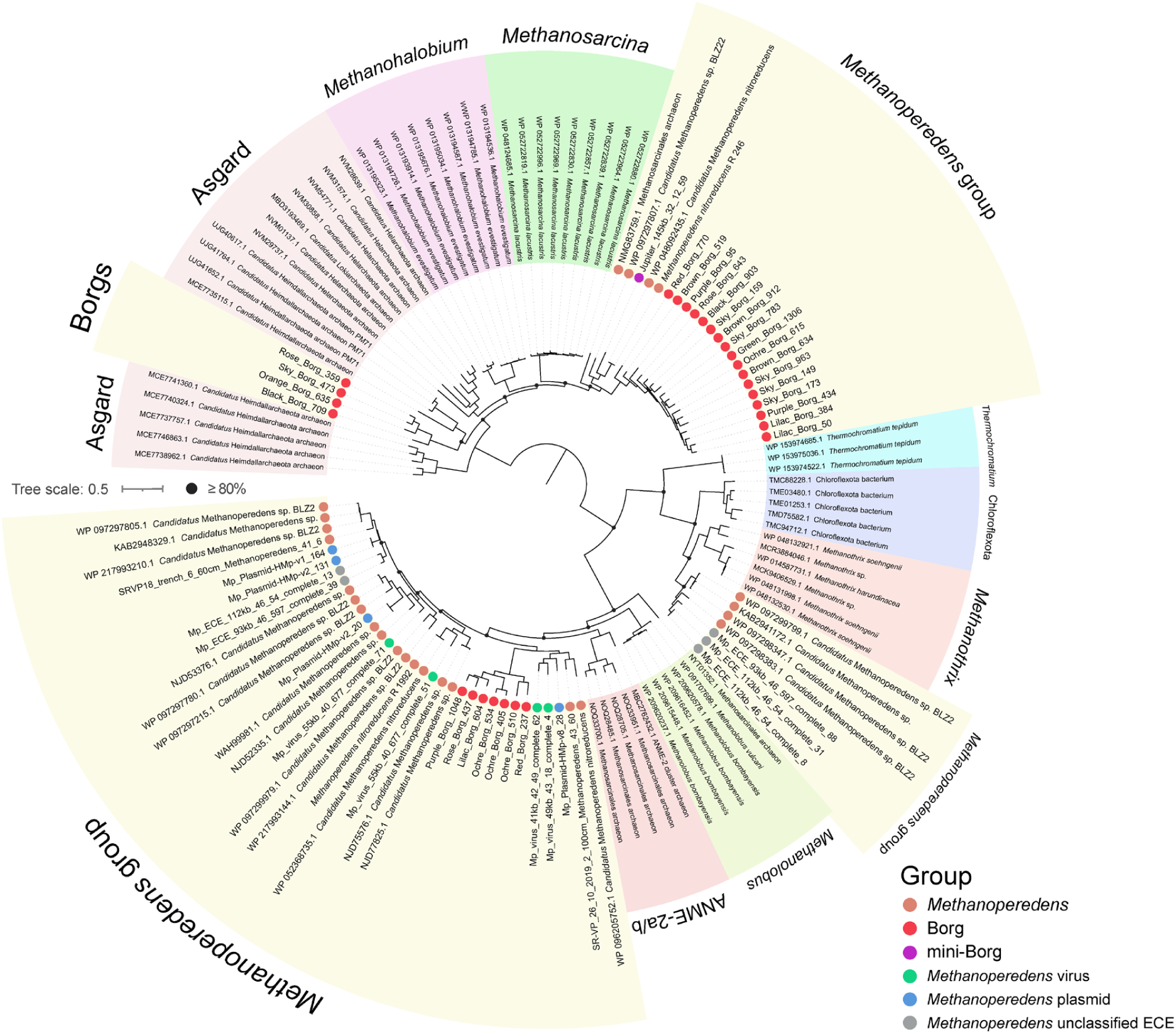
Phylogeny of TnpB transposase shared by *Methanoperedens* and associated ECEs. Homologous references were recruited from the NCBI nr database. Arc colors indicate different taxonomic clades. Proteins found in *Methanoperedens* and associated ECEs are marked using colored dots. The tree was mid-point rerooted and support values were calculated based on 1000 replicates.

**Fig. S22.**
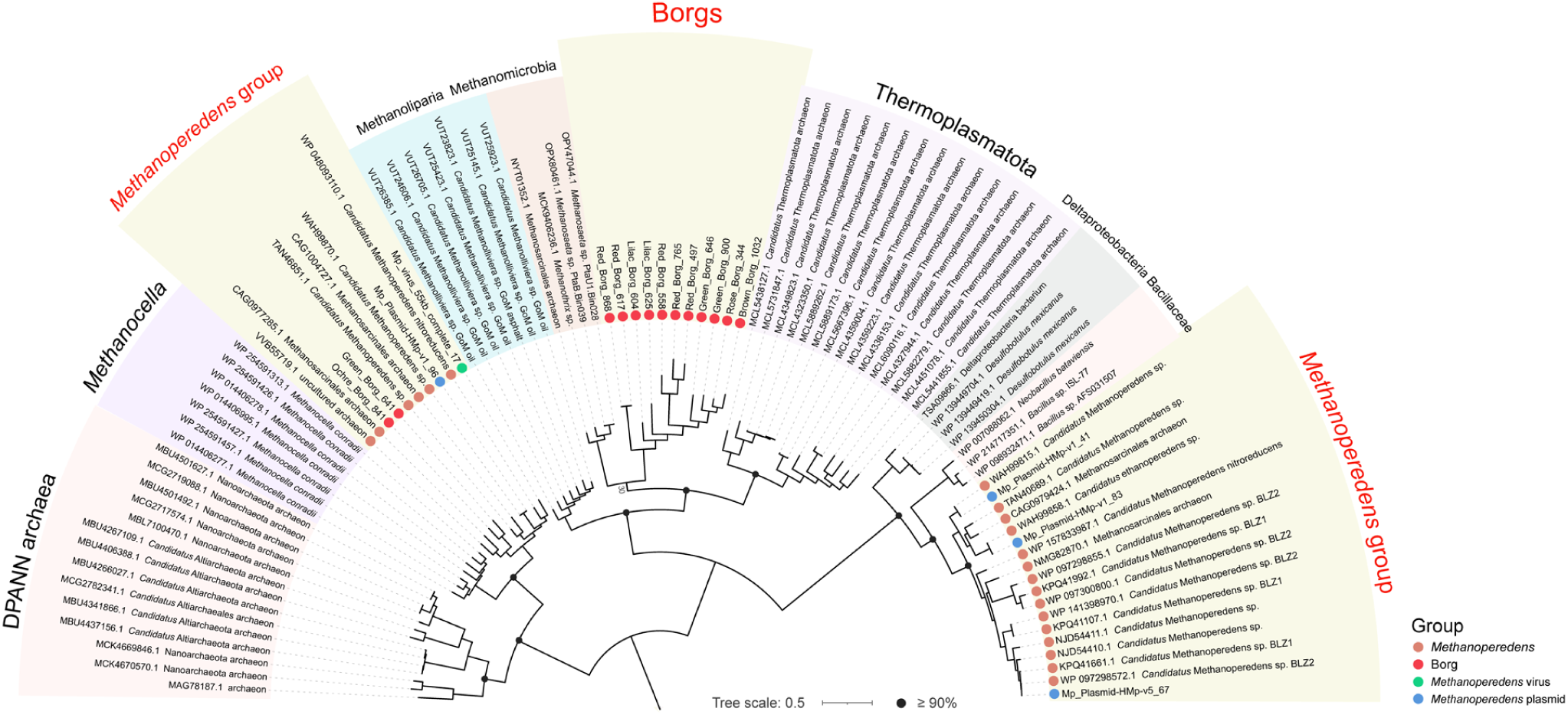
Phylogeny of non-TnpB transposases shared by *Methanoperedens* and associated ECEs. Homologs were recruited from the NCBI nr database. Arc colors indicate different taxonomic clades. Proteins found in *Methanoperedens* and associated ECEs are marked using colored dots. The tree was mid-point rerooted and support values were calculated based on 1000 replicates.

**Fig. S23.**
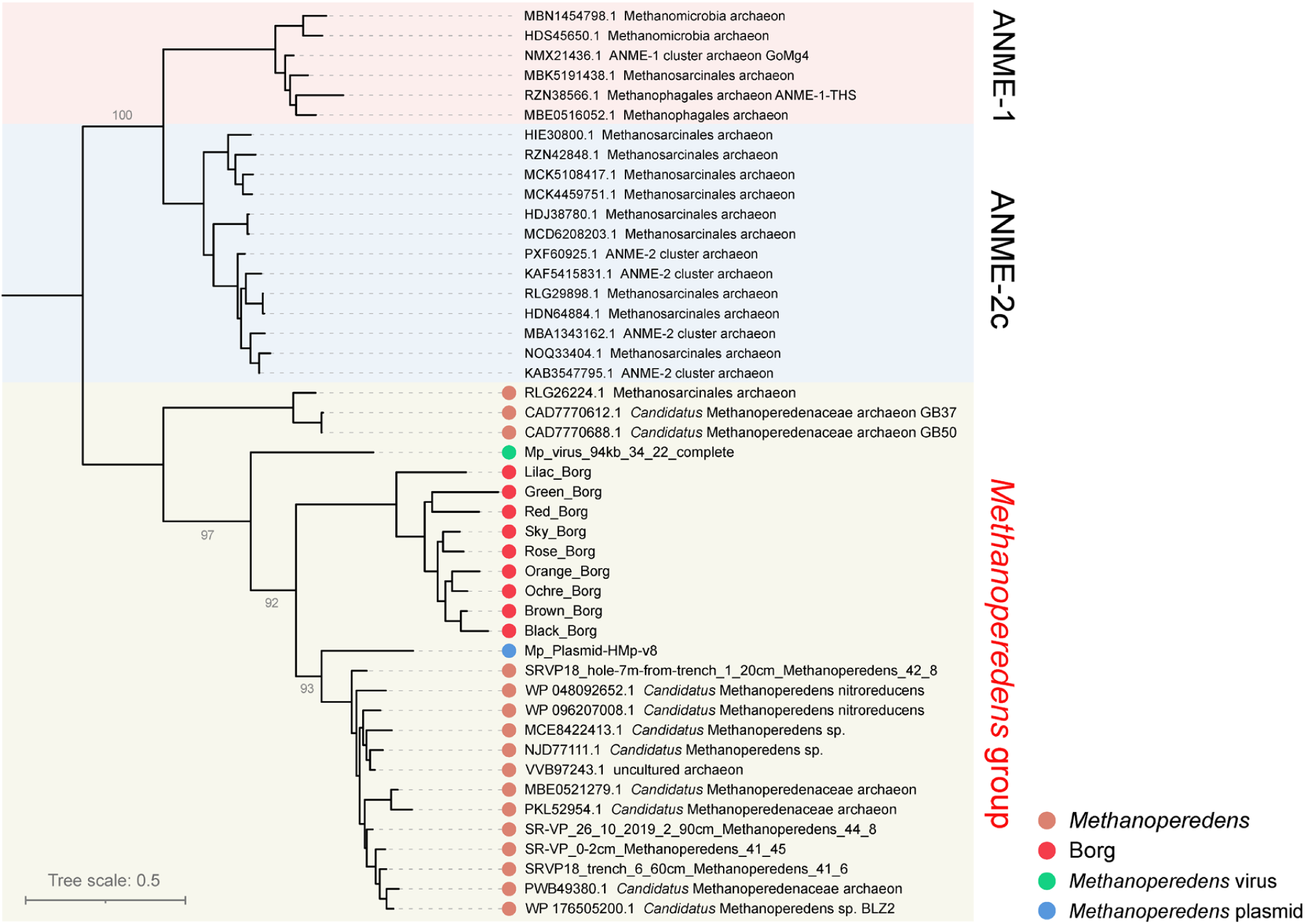
Phylogeny of classical thymidylate synthase (ThyA) shared by *Methanoperedens* and associated ECEs. Homologous references were recruited from NCBI nr. Proteins found in *Methanoperedens* and associated ECEs are marked using colored dots. Support values were calculated based on 1000 replicates.

**Fig. S24.**
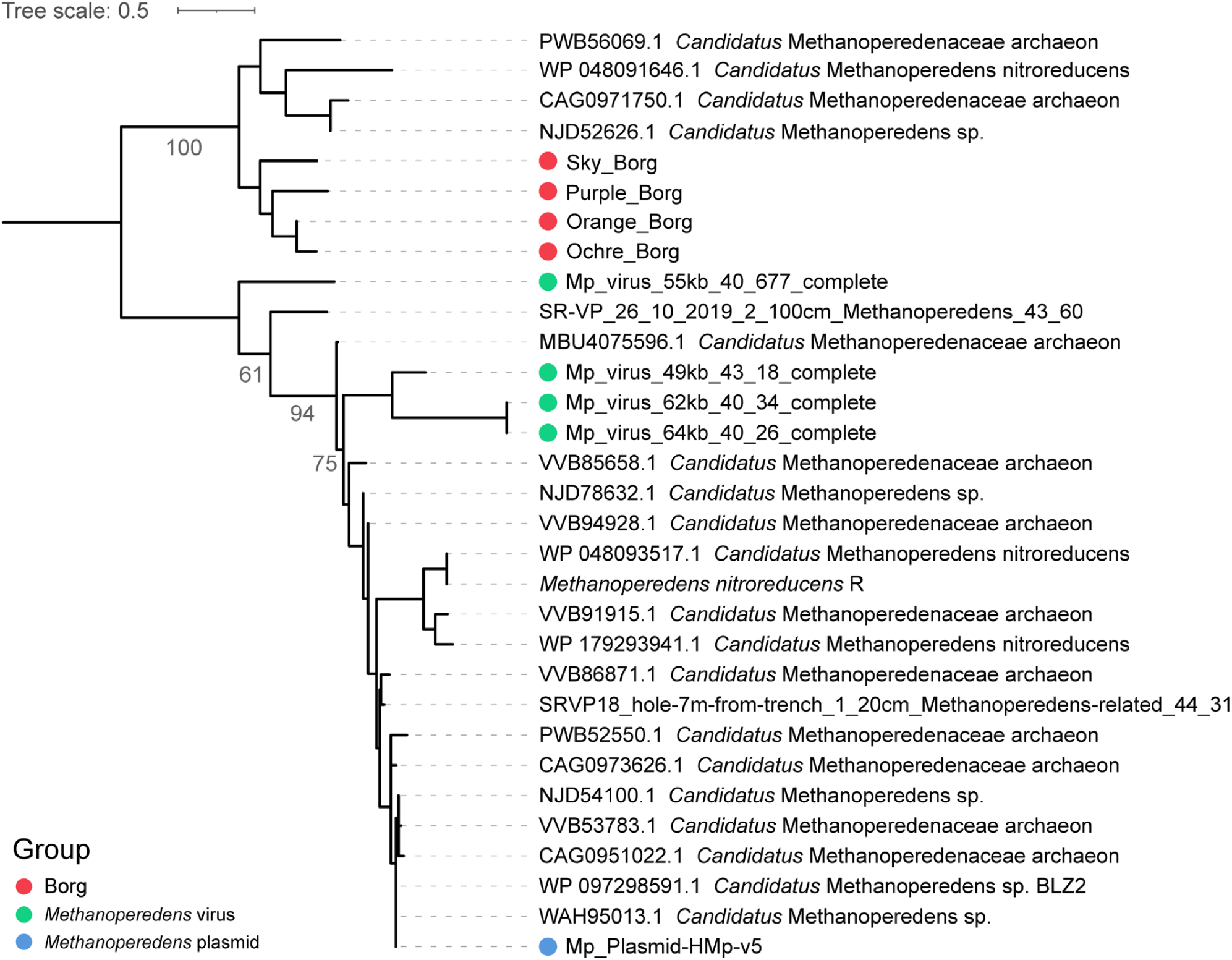
Phylogeny of a hypothetical protein (subfam2653) shared by *Methanoperedens* and associated ECEs. The only similar proteins identified in NCBI nr belong to *Methanoperedens*. The tree was mid-point rerooted and support values were calculated based on 1000 replicates.

**Fig. S25.**
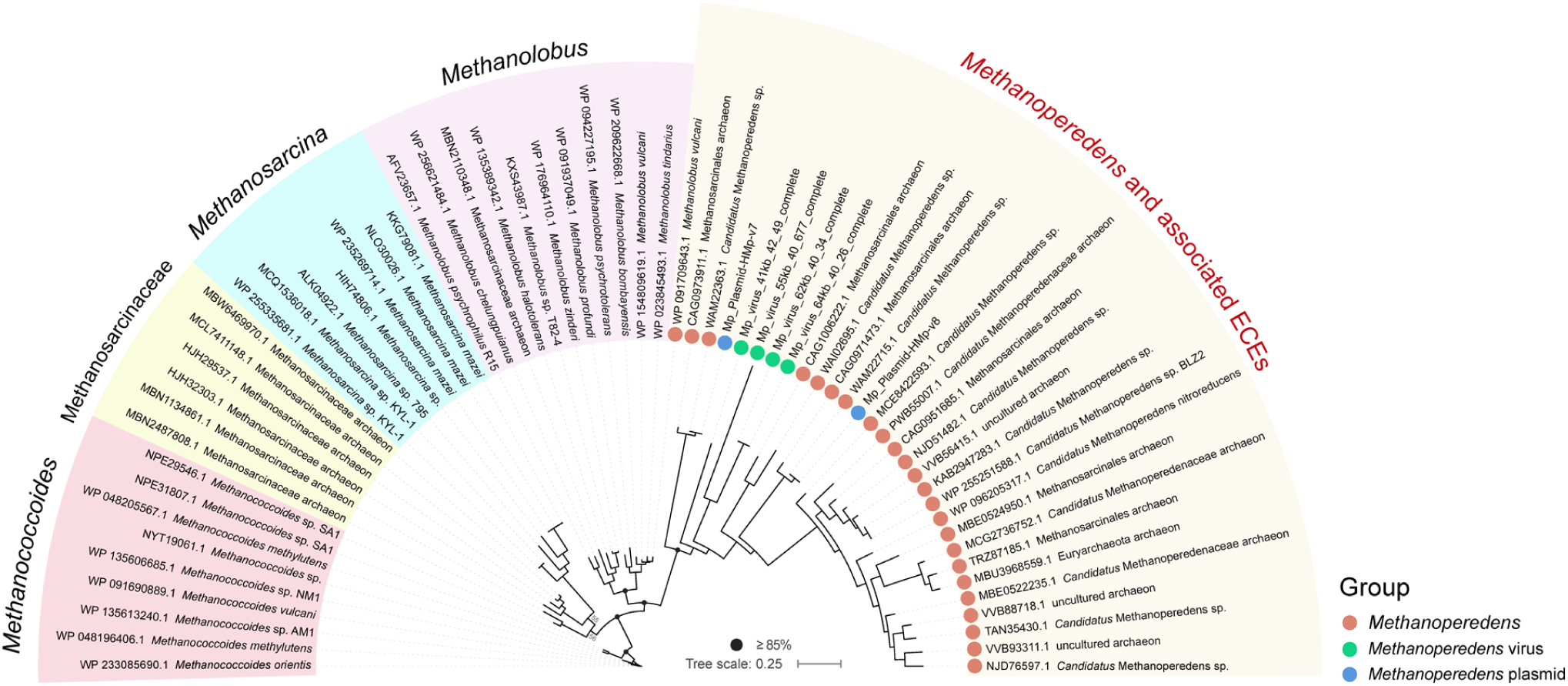
Phylogeny of DNA repair endonuclease (ERCC4) shared by *Methanoperedens* and associated ECEs. Homologous references were recruited from NCBI nr. Arc colors indicate different taxonomic clades. Proteins found in *Methanoperedens* and associated ECEs are marked using colored dots. The tree was mid-point rerooted and support values were calculated based on 1000 replicate.

**Table S1.** Genomic features of mini-Borg genomes.

| Mini-Borg genomes | Genome size (bp) | GC content | #ORFs | ITR length (if there) | #TR regions | #TR in ORFs | #Intergenic TR | #Replichores |
| --- | --- | --- | --- | --- | --- | --- | --- | --- |
| Venus_106kb_34_23_complete | 106,520 | 33.9% | 145 | 223 bp | 15 | 10 | 5 | 2 |
| Saturn_96kb_32_15_complete | 96,760 | 31.8% | 130 | 52 bp | 9 | 6 | 3 | 2 |
| Saturn_91kb_33_14_complete | 91,043 | 33.0% | 129 | 166 bp | 4 | 4 | 0 | 2 |
| Saturn_82kb_33_70_complete | 82,145 | 32.9% | 113 | 85 bp | 4 | 1 | 3 | 2 |
| Jupiter_60kb_34_58_complete | 60,376 | 34.3% | 76 | 669 bp | 10 | 6 | 4 | 1 |
| Uranus_55kb_35_16_complete | 55,475 | 34.9% | 83 | 118 bp | 8 | 2 | 6 | 1 |
| Jupiter_54kb_36_306_complete | 54,669 | 35.6% | 72 | 237 bp | 5 | 4 | 1 | 1 |
| Jupiter_52kb_35_101_complete | 52,459 | 34.5% | 70 | 25 bp | 0 | 0 | 0 | 1 |
| Jupiter_144kb_32_101_near_complete | 144,326 | 32.2% | 206 | 363 bp | 3 | 3 | 0 | ND |
| Saturn_94kb_33_10_near_complete | 93,177 | 32.5% | 119 | NA | 1 | 0 | 1 | 2 |
| Saturn_75kb_34_20_near_complete | 75,189 | 34.1% | 103 | NA | 0 | 0 | 0 | 2 |
| Jupiter_60kb_33_14_near_complete | 60,241 | 32.7% | 81 | NA | 11 | 5 | 6 | 2 |
| Jupiter_145kb_32_12 | 145,330 | 32.0% | 197 | NA | 11 | 6 | 5 | ND |
| Jupiter_112kb_33_38 | 112,613 | 32.9% | 171 | NA | 3 | 1 | 2 | ND |
| Neptune_111kb_32_42 | 111,664 | 32.0% | 164 | NA | 8 | 4 | 4 | 2 |
| Jupiter_94kb_32_17 | 94,633 | 32.4% | 113 | NA | 7 | 2 | 5 | ND |
| Jupiter_83kb_35_5 | 83,085 | 34.2% | 112 | NA | 2 | 1 | 1 | 2 |
| Uranus_69kb_33_33 | 69,134 | 32.7% | 103 | NA | 2 | 2 | 0 | ND |
| Jupiter_55kb_35_19 | 55,519 | 34.6% | 80 | NA | 2 | 1 | 1 | ND |
| Jupiter_54kb_36_83 | 54,901 | 35.9% | 79 | NA | 1 | 1 | 0 | ND |
| Neptune_46kb_32_14 | 46,651 | 31.9% | 72 | NA | 0 | 0 | 0 | 2 |
| Saturn_41kb_33_22 | 41,779 | 32.9% | 77 | NA | 5 | 2 | 3 | ND |
| Jupiter_39kb_32_16 | 39,341 | 31.6% | 62 | NA | 6 | 3 | 3 | ND |

Note: NA means "not available"; ND means "not determined".

**Table S2.**
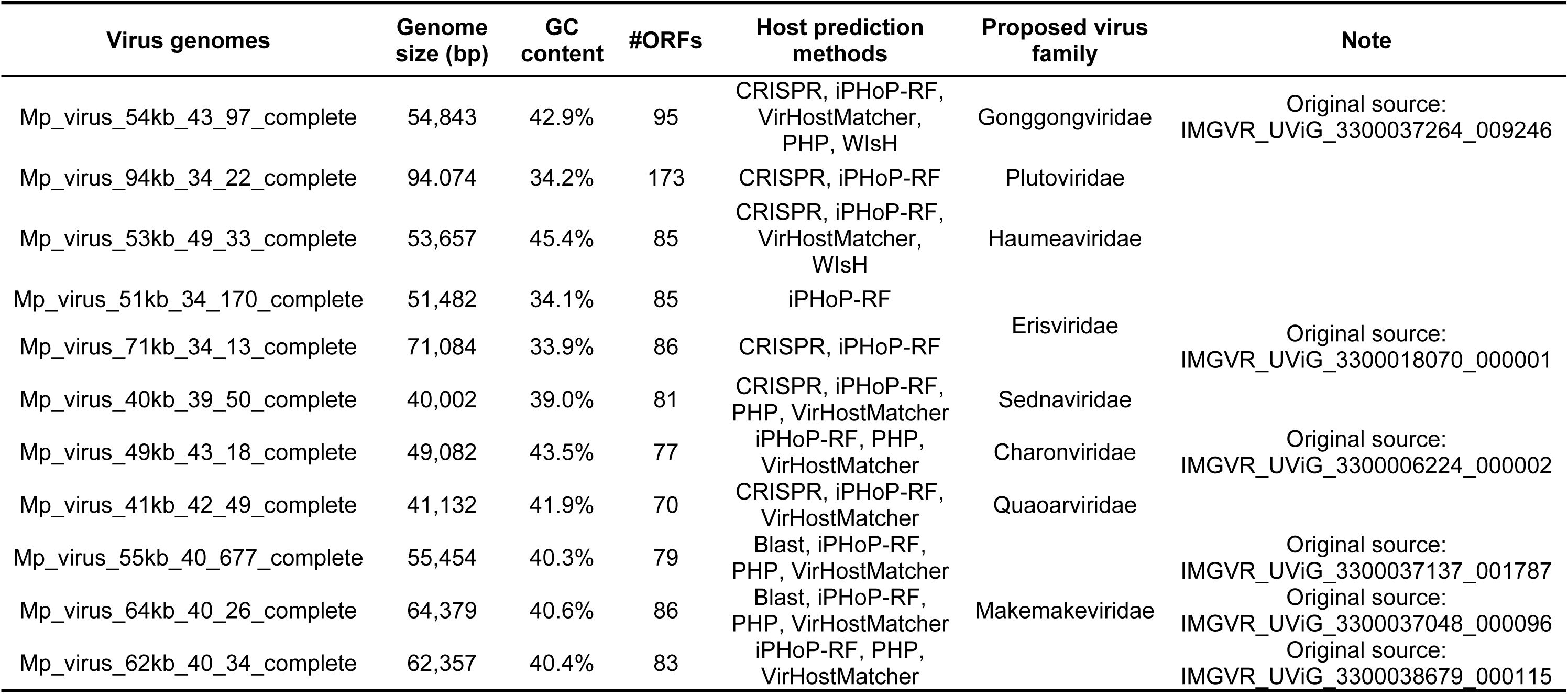
Genomic features of viral genomes.

**Table S3.** Genomic features of unclassified ECEs.

| Unclassified ECE genomes | Genome size (bp) | GC content | #ORFs | Host prediction methods |
| --- | --- | --- | --- | --- |
| Mp_ECE_93kb_46_597_complete | 93,558 | 46.0% | 148 | CRISPR, Blast |
| Mp_ECE_112kb_46_54_complete | 112,435 | 45.4% | 165 | CRISPR |
| Mp_ECE_66kb_47_24_complete | 66,699 | 46.6% | 105 | Blast |
| Mp_ECE_14kb_44_38_complete | 14,272 | 43.9% | 21 | Blast |

